# HEIP1 orchestrates pro-crossover protein activity during mammalian meiosis

**DOI:** 10.1101/2025.08.25.672081

**Authors:** Arnaud De Muyt, Sunkyung Lee, Sushil Khanal, Laurine Dal Toe, Céline Adam, Raphael Mercier, Valérie Borde, Neil Hunter, Thomas Robert

## Abstract

Meiotic crossovers are needed to produce genetically balanced gametes. In mammals, crossover formation is mediated by a conserved set of pro-crossover proteins via mechanisms that remain unclear. Here, we characterize a mammalian pro-crossover factor HEIP1. In mouse HEIP1 is essential for crossing over and fertility of both sexes. HEIP1 promotes crossing over by orchestrating the recruitment of other pro-crossover proteins, including the MutS*γ* complex (MSH4- MSH5) and E3 ligases (HEI10, RNF212, and RNF212B), that are required to mature crossover sites and recruit the crossover-specific resolution complex MutL*γ*. Moreover, HEIP1 directly interacts with HEI10, suggesting a direct role in controlling the recruitment of pro-crossover E3 ligases. During early stages of meiotic prophase I, HEIP1 interacts with the chromosome axes, independently of recombination, before relocalizing to the central region of the synaptonemal complex. We propose that HEIP1 is a new conserved master regulator of crossover proteins that controls different crossover maturation steps.

**Significance Statement:** Crossovers are essential to produce gametes by promoting the proper segregation of the homologous chromosomes. But, if misregulated, they can lead to genetic disorders, miscarriage and infertility. Their formation depends on the conserved pro-crossover factors, which repertoire is expanding and mode of action tightly regulated. This study highlights how the HEIP1 protein organizes pro-crossover protein activities in the mouse. Our findings show that HEIP1, by interacting early with chromosomes independently of recombination initiation, and orchestrating the recruitment of pro-crossover factors, including the MutSg complex and the RING proteins HEI10, RNF212 and RNF212B, is pivotal in this regulation. Our work is of significance to unravel crossover control during meiotic recombination, a conserved mechanism essential for gametes formation.

## Introduction

The specialized meiotic program, separates homologous chromosomes at the first meiotic division (meiosis I) to enable ploidy reduction in the gametes needed for sexual reproduction (1). In most species, the formation of a physical connection between homologs, the chiasma, is essential for their proper segregation. Chiasmata are the cytological correlates of the DNA crossovers (COs) that arise from the resolution of homologous recombination (HR) intermediates formed in meiotic prophase I (1, 2). In meiosis, HR is initiated by the programmed formation of DNA double strand breaks (DSBs) that are catalyzed by the topoisomerase-like complex TOPOVIL (SPO11-TOPOVIBL) and its accessory factors (3–6). Following DSB formation, removal of SPO11 from DSB ends, resection of the 5’ strands, and 3’-strand invasion of a homologous duplex, catalyzed by the DMC1/RAD51 strand exchange proteins, lead to the formation of nascent displacement loops (D-loop) (1). A subset of D-loops will be stabilized into single-end invasions, converted into double Holliday junctions (dHJ) and finally resolved into COs through one of two pathways that define class I and II events, which co-exist in various organisms (7). In the class I pathway, the final resolution of dHJs is catalyzed by the MutL*γ* endonuclease complex, composed of the MLH1 and MLH3 proteins, and leads exclusively to COs (8–11). In the mouse, MLH1 and MLH3 colocalize as discrete foci on the meiotic chromosome axis, from mid to late pachynema. Generally, one-two foci per bivalent are detected that colocalize with the multiprotein recombination nodules that mark COs and chiasma sites (12–16). In *Mlh1^-/-^* and *Mlh3^-/-^*mice, recombination nodules disappear, and chiasma formation and COs are diminished (12, 14, 17). In the mouse, 90% of COs are formed through the class I pathway (9, 18). The class II pathway depends on the resolution of joint molecules by structure-specific endonucleases, including MUS81-EME1 and SLX1. The majority of recombination events do not mature into crossovers and are instead resolved as non-crossovers by the Bloom complex (BLM-TOP3ɑ-RMI1-RMI2) (11, 19, 20).

Class I COs depend on a well conserved set of meiotic pro-CO proteins, originally identified in *Saccharomyces cerevisiae* as the ZMM proteins (Zip1-4, Msh4, Msh5, Mer3 and Spo16) (7). In budding yeast, these proteins form subcomplexes that interact with DNA recombination intermediates and are thought to work collectively to promote and protect dHJs, and direct their resolution by the MutL*γ* complex toward COs. Class I COs are subject to interference, a process that prevents the formation of two COs in close vicinity (21, 22). The molecular mechanisms underlying the formation of class I COs and interference remain unclear. In many species, Zip3- related proteins initially localize as numerous discrete foci along the synaptonemal complex (SC), the proteinaceous structure that assembles between synapsed chromosomes, before progressively concentrating into a few larger brighter foci that colocalize with MLH1 and mark the CO sites (23–27). In *Arabidopsis thaliana*, the dynamics of the Zip3-family protein HEI10 can be modeled by a SC-dependent coarsening diffusion process that leads to the interference patterning of class I CO sites (28–30). However, it is unclear how HEI10 is initially recruited to prophase chromosomes and how it cooperates with other pro-CO components. Two recent studies performed in rice and *Arabidopsis* identified a direct interactor of HEI10 named HEIP1 (for HEI10 Interacting Protein) as a new candidate ZMM factor required to regulate HEI10 and the patterning of class I CO sites (31, 32). In rice, *heip1* mutations cause diminished HEI10 foci at late pachynema and reduce crossing over (31). Similarly, in *A. thaliana*, *heip1* mutants are defective for crossing over in the class-I pathway (32). At the molecular level, *A. thaliana* HEIP1 was inferred to exert a dual role, upstream of MLH1 to control HEI10 accumulation, but also downstream, to promote CO maturation into chiasma (32). Importantly, HEIP1 was shown to be conserved, with potential orthologues in mammals, suggesting a conserved function in regulating HEI10 (32).

In mammals, the ZMM homologs are SYCP1(Zip1), SHOC1 (Zip2), TEX11 (Zip4), MSH4- MSH5, HFM1 (Mer3), and SPO16 plus three Zip3-related proteins RNF212, RNF212B, and HEI10 (7, 33, 34). The three pro-crossover RING-domain proteins (CORs) are inferred to define two subgroups, RNF212/RNF212B and HEI10, that control the stability of other ZMMs through SUMO and ubiquitin modifications, respectively (7, 26). In the absence of RNF212, COs are reduced by more than 90%, MLH1/MLH3 foci are not detected, and the turnover of MSH4/MSH5 foci is accelerated. Therefore, RNF212 might contribute to the stabilization of ZMM proteins and recombination intermediates to mediate CO site designation. HEI10 is a putative E3 ubiquitin ligase that is also essential for crossing over (35). In *Hei10^-/-^* mice, initial formation of MSH4/MSH5 foci is normal but high numbers of foci then persist throughout pachynema and MLH1/MLH3 foci, marking designated CO sites, fail to form (26, 35). These results suggest that HEI10 promotes CO-site designation and maturation. It has been suggested that RNF212 and HEI10 form a SUMO/ubiquitin relay that regulates MutS*γ*-dependent stabilization of single-end invasions. However, the precise mode of action of each COR, their targets and partners remain unknown. An additional layer of complexity was recently added by the identification of RNF212B, a third mammalian COR that is also essential for the formation MLH1 foci and COs, and acts non-redundantly with HEI10 and RNF212 (33, 34). Thus, CORs appear pivotal for CO designation and maturation, but unraveling their regulation remains a priority to fully understand the control of class I COs in mammals. Particularly, deciphering their mode of recruitment and interplay with recombination and SC components is of crucial interest to evaluate models of crossover patterning such as the SC-dependent coarsening process (29), or physical tension proposed by the beam-field model (36).

Two studies of a human gene implicated in nonobstructive azoospermia identified the mouse *Redic1* protein as a class I CO factor (37, 38). Point mutations in *Redic1* cause defective spermatogenesis, diminished crossing over and strong reductions in the assembly of MutS*γ* and MutL*γ* foci. Phylogenetic analysis suggested that the *Redic1* sequence corresponds to the *A. thaliana Heip1* sequence (32). However, it remains unclear whether *M. musculus* REDIC1/HEIP1 is a functional orthologue of *A. thaliana* HEIP1.

Here, we perform extensive characterization of mouse HEIP1. We show that mouse HEIP1 is essential for fertility, and promotes crossing over by regulating the ZMMs: MSH4, TEX11, MLH1 and the E3 ligases HEI10, RNF212, and RN212B. At the molecular level, HEIP1 is initially loaded onto meiotic chromosome axes before they synapse and independently of HR. It then relocalizes between the closely juxtaposed axes, but independently of the SC central element proteins, following RPA ssDNA binding complex. These observations suggests that HEIP1 acts at an early stage of meiotic recombination to stabilize intermediates and promote CO-site designation. The number of HEIP1 foci decreases throughout pachynema and then it colocalizes specifically to crossover sites marked by MLH1, indicating that HEIP1 also has a later role in CO maturation. Moreover, HEIP1 directly interacts with HEI10, and is required for its recruitment to CO sites. RNF212/RNF212B foci aberrantly accumulate in *Heip1^-/-^* spermatocytes, suggesting that HEIP1 plays a pivotal role in orchestrating the COR-mediated SUMO/ubiquitin relay. Altogether, we propose that HEIP1 is a key regulator of class-I CO designation and maturation, exerting a dual function through interaction with the chromosome axes and controlling the recruitment and stabilization of ZMM factors.

## Results

### HEIP1 is expressed predominantly during meiotic prophase I and is essential for fertility

Human C12ORF40 was recently identified as a putative homolog of *A. thaliana* HEIP1 (32). C12ORF40/HEIP1 (hereafter called HEIP1) is a protein of 652 amino acids predicted to be predominantly disordered, except for tandem *α*-helices at its N-terminus, suggesting it may behave like an intrinsically disordered protein (Figure S1A). To evaluate HEIP1 function in mouse, we first monitored its expression. *Heip1* was specifically expressed in testes, but not in somatic tissues (Figure S1B). Temporal analysis of *Heip1* expression, during the first synchronous wave of spermatogenesis in testes from juvenile males, revealed a peak of expression at 12-14 days post- partum (dpp), during early meiotic prophase I (Figure S1C), compatible with a role in meiotic recombination.

To investigate HEIP1 function, a constitutive *Heip1* knockout mouse line was engineered by CRISPR-Cas9 DSB targeting, to generate a 4 bp deletion that introduces a frameshift resulting in the replacement of K43 by five amino acids (SPLSV) followed by a stop codon (Figure S1A, and see Figure S1D, Material and methods for genotyping strategy). This mutation is predicted to result in a severely truncated HEIP1 protein (Figure 1A). In *Heip1^−/−^* mice, HEIP1 protein was undetectable by immunocytochemistry of surface-spread spermatocyte chromosomes (Figure S1E). *Heip1^−/−^* mice developed normally and reached adulthood but both males and females were sterile (no pups born in 6 months from 6 crosses each of *Heip1^-/-^* males and females with wild type C57/BL6). *Heip1^−/−^* testes were ∼3 times smaller than those of wild type animals (Figure 1B) and histological analysis showed the absence of post-anaphase I cells, suggesting spermatocyte depletion at this stage (Figure 1C). Meiotic arrest at anaphase I is frequently observed in CO defective mutants (*Rnf212^−/−^*, *Hei10^−/−^*, and *Mlh3^−/−^*) (14, 35, 39), while meiotic arrest at the pachytene stage of prophase I is detected in mutants defective for meiotic DSB formation and/or repair (*Spo11^−/−^*, *Dmc1^−/−^*) or synapsis (*Sycp1^−/−^*) (40, 41). Thus, CO may be impaired in *Heip1^−/−^*spermatocytes. Consistent with a lack of chiasmata in *Heip1^−/−^*, chromosome congression was severely defective in metaphase I spermatocytes (Figure 1D, arrows). Histological analysis of ovaries showed that follicle number was strongly reduced in *Heip1^−/−^* females compared with wild type animals, consistent with an essential role for HEIP1 in oogenesis (Figure S1F).

**Figure 1.**
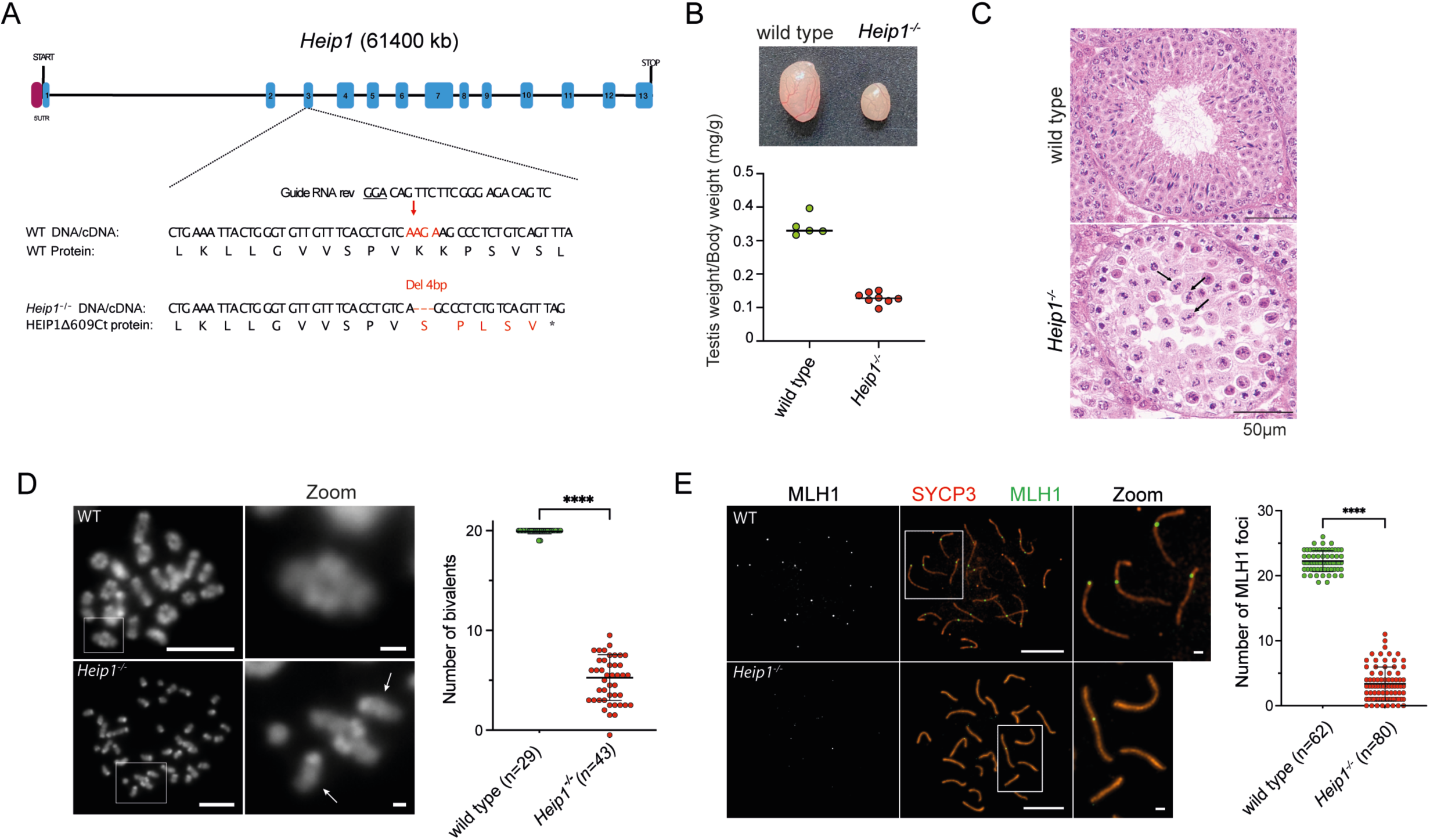
*Heip1* deletion leads to massive germline loss and infertility. (A) Schematic representation of the *Heip1* locus showing exons (blue boxes), untranslated regions (5′-UTR, purple box), and the start of the open reading frame in exon 1. View of exon 3 shows the guide RNA sequences (guide RNA rev with PAM underlined) used for CRISPR/Cas9 editing, the genomic DNA/cDNA sequence of the edited region and the corresponding wild type (WT) and *Heip1^-/-^* protein sequences. DNA break sites are marked by red arrows; red text color marks the mutated DNA and protein sequences. (B) Representative image of wild type and *Heip1^-/-^* testes (top). Testis to body weight ratios of 2-month-old wild type and *Heip1^-/-^* mice. Each dot corresponds to the mean value for the two testes from one mouse. Data are the mean ± SD. ****P < 0.0001 (two-tailed Student’s *t*-test). (C) Representative images of seminiferous tubules stained with hematoxylin and eosin from testis sections of 60 dpp wild type and *Heip1^−/−^* mice. In *Heip1^−/−^* seminiferous tubules, cells progress to metaphase I (stage XII) (see arrows), normal chromosome congression is not observed, and post-meiotic spermatogenic cells (spermatids and spermatozoa) are absent. Scale bar, 50 μm (D) Left: DAPI staining of diakinesis/metaphase I chromosome spreads and magnified images from wild type and *Heip1^-/-^* spermatocytes from 60 dpp mice. Scale bar, 10 μm. Arrow indicates a univalent in a *Heip1^−/−^* cell. Scale bar, 1 μm. Right: number of bivalents per nucleus Each dot corresponds to one nucleus. (E) Immunostaining of MLH1 and SYCP3 on chromosome spreads from wild type and *Heip1^-/-^* spermatocytes at 18 dpp. White squares correspond to the zoom images on the right. The number of MLH1 foci is presented on the right. Scale bar, 10 µm. For all graphs, ****P < 0.0001 (two-tailed unpaired Mann-Whitney test).

### Crossover formation is reduced in *Heip1^-/-^* spermatocytes

To directly test the role of HEIP1 in CO formation, we first determined whether homologs were connected as bivalents by chiasmata, the cytological correlates of COs, in metaphase-I spermatocyte chromosome spreads (Figure 1D). In wild-type cells, the number of bivalents was twenty compared to only five in *Heip1^-/-^* cells, confirming diminished crossing over. To confirm that the class I CO pathway was specifically impaired in the *Heip1^-/-^* mutant, spermatocyte chromosome spreads were immunostained for the class I CO markers MLH1 and CNTD1, a meiosis specific cyclin-homolog that acts in concert with MutLg (42) (Figure 1E and Figure S1G). The mean numbers of both MLH1 and CNTD1 foci was 22±2 in wild-type spreads, but was decreased by 85% (MLH1) and 84% (CNTD1) in *Heip1^-/-^*spermatocytes, indicating that HEIP1 is required for interference-dependent class I CO formation.

### HEIP1 controls the stabilization of mid-stages recombination intermediates, but not the early steps of meiotic HR

To identify the step(s) of recombination controlled by HEIP1, the efficiency of DSB formation was assessed *via* immunolocalization of MEI4, an essential component of the DSB machinery (43), and H2AX phosphorylation (γH2AX), a marker of DSB formation (Figure S2A-B). In wild type spermatocytes, numerous MEI4 foci were observed along the chromosome axes at leptonema; their number decreased as synapsis progressed during zygonema, and most foci were removed by pachynema (Figure S2A). γH2AX was present over large chromatin domains during leptonema and zygonema and specifically accumulated on the sex chromosomes during pachynema forming the sex body (Figure S2B). In *Heip1^-/-^* spermatocytes, immunolocalization of MEI4 and γH2AX was similar to that observed in wild type spermatocytes (Figure S2A-B), suggesting that HEIP1 is not required to recruit the DSB machinery to chromosome axes and form DSBs. DSB end processing and strand invasion steps were monitored via immuno-localization of RPA2, SPATA22 and DMC1 (Figure 2A-B and Figure S2C). RPA2 and SPATA22 are components of the RPA and SPAT22-MEIOB complexes that bind single-stranded DNA (ssDNA) generated at resected DSBs and D-loops following strand exchange (44). The meiosis-specific RAD51 homolog DMC1 assembles onto resected DSB ends to form into nucleoprotein filaments that catalyze homology search and strand exchange reactions. In wild-type spermatocytes, RPA2, SPATA22 and DMC1 form numerous foci on the chromosome axes from leptonema to zygonema that then decrease at pachynema (Figure 2A-B and Figure S2C). In *Heip1^-/-^* spermatocytes, the dynamic localization patterns of DMC1, RPA2 and SPATA22 were unaffected, indicating that HEIP1 is not required for the early HR steps of DNA end processing and single-strand invasion (Figure 2A-B and Figure S2C). Consistently, we did not observe any defect in prophase I progression in *Heip1^-/-^*spermatocytes (Figure S2D).

**Figure 2.**
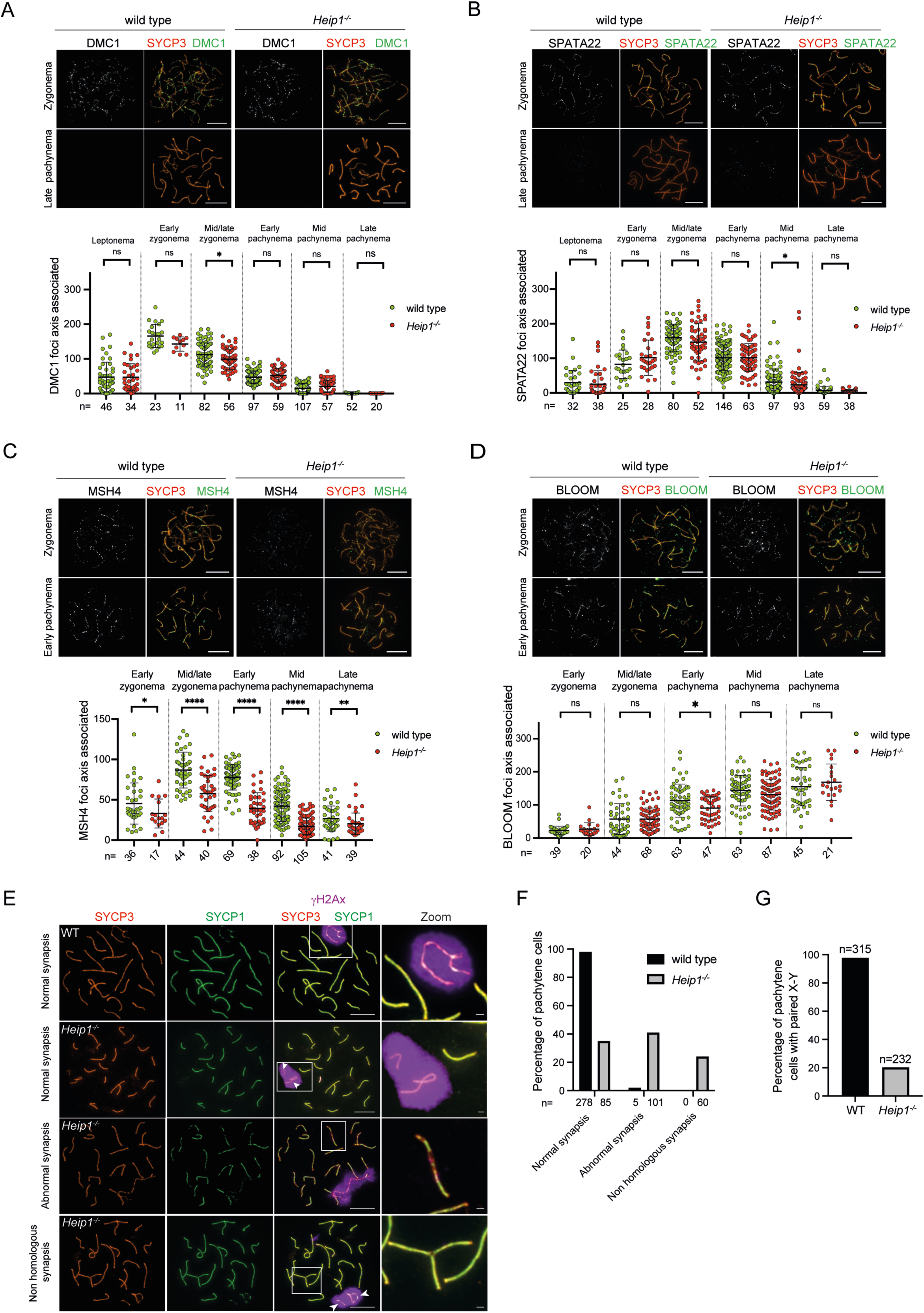
Homologous recombination and synapsis in wild type and *Heip1^-/-^* mice. (A-D). Analysis of homologous recombination in wild type and *Heip1^-/-^* mice. (A) SYCP3 (orange) and DMC1 (green), (B) SYCP3 (orange) and SPATA22 (green), (C) SYCP3 (orange) and MSH4 (green), and (D) SYCP3 (orange) and Bloom (green) localization in 18 dpp wild type and *Heip1^-/-^* spermatocytes at different prophase I substages. Scale bar, 10 μm (all panels). Plots at the bottom of the panels show the quantification of foci; ns, not significant, *P < 0.1, **P < 0.01, ****P < 0.0001 (two-tailed unpaired Mann-Whitney test). (E) Chromosome synapsis is defective in *Heip1^-/-^* mice. Representative images of chromosome spreads from 60 dpp wild type and *Heip1^-/-^* spermatocytes immunostained for ɣH2Ax (purple), SYCP3 (red) and SYCP1 (green). Magnified images from the white boxes show representative chromosomes. Scale bars, 10 μm for full nuclei and 1 μm for the magnified images. (F) Percentages of pachytene spermatocytes with and without synapsis defects in wild type and *Heip1^-/-^* mice. Cells were divided into three groups: with fully synapsed chromosomes (normal synapsis); with discontinued or unsynapsed chromosomes (abnormal synapsis); with synapsis between non-homologous chromosomes (non-homologous synapsis). Experiments were performed at least twice and one experiment is shown. (G) Percentages of pachytene spermatocytes with paired X-Y chromosomes in wild type and *Heip1^-/-^* mice.

Post strand invasion steps of crossing over are promoted by ZMM-related factors, that include the MutS*γ* complex, MSH4-MSH5. Immunofluorescence staining of MSH4 in wild-type spermatocytes revealed initial localization to synapsed regions during zygonema, with focus numbers reaching a maximum mean of 98±21 foci (Figure 2C). In *Heip1^-/-^*spermatocytes, MSH4 foci numbers were reduced by 33% in late zygonema and then by ≥50% in early/mid pachynema, suggesting a defect in the maintenance of MutS*γ* complexes required to implement crossing over. We also examined the localization dynamics of a second ZMM pro-crossover factor, TEX11 (Figure S2E)(45, 46). Similar to MSH4, in wild-type spermatocytes, TEX11 localized to synapsed chromosomes and reaching a peak of 118±35 foci in early pachynema. Then, TEX11 focus numbers rapidly decreased after early pachynema and almost completely disappeared by mid pachynema (Figure S2E). In *Heip1^-/-^* spermatocytes, TEX11 foci were reduced by 85% in early pachynema (Figure S2E), indicating that HEIP1 is also required to stabilize TEX11 foci.

In budding yeast *zmm* mutants, recombination intermediates are destabilized by an anti-crossover activity of Sgs1-Top3-Rmi1, ortholog of the mammalian Bloom complex (BLM-TOPIIIa-RMI1/2), resulting in non-crossover formation (47, 48). Therefore, localization of the Bloom helicase BLM was examined to determine whether HEIP1 affects its loading. In wild-type spermatocytes, BLM localized first as numerous foci on synapsed chromosomes during zygonema and reached a plateau as synapsis was completed in pachynema (Figure 2D). As expected, we noticed that the BLOOM protein is not recruited in *Top6bl^-/-^* mutants, confirming that the loading of the protein depends on recombination initiation (Figure S2F). In *Heip1^-/-^*spermatocytes, numbers of BLM foci remained largely unchanged, suggesting that HEIP1 does not regulate BLM dynamics.

Altogether, these data suggest that HEIP1 is required to stabilize pro-CO factors, represented here by MSH4 and TEX11, but does not appear to influence early steps of meiotic HR including DSB formation and strand invasion nor the chromosomal dynamics of the Bloom helicase.

### Autosomal and X-Y synapsis are impaired in *Heip1*^-/-^ spermatocytes

In mice, mutants defective in DSB formation or repair are typically defective for homolog synapsis, leading to pachytene arrest and apoptosis of spermatocytes (5, 43, 49–51). However, among mutants that affect later steps of meiotic recombination, the nature of synapsis defects is more variable. For instance, *Hei10^mei4/mei4^*, *Mlh1^−/−^*, *Mlh3^−/−^*, *Prr19^−/−^* and *Cntd1^−/−^* mice (CO mutants) do not show overt SC defects (14, 17, 35, 42, 52) nor meiotic arrest, while *Shoc1^−/−^*, *Spo16^−/−^* and *Msh4^−/−^* mice show aberrant synapsis, defective XY sex-body formation, and arrest at pachytene (53–55). Finally, *Hfm1^−/−^*, *Rnf212^−/−^, and Rnf212b^−/−^*mice show modest SC defects but no arrest at pachytene (35, 39, 56). To monitor synapsis and formation of the XY sex body in *Heip1^-/-^*mutant spermatocytes, chromosome spreads were immunostained for the axis protein SYCP3, transverse filament SYCP1, and γH2AX (Figure 2E). Only 35% of pachytene-like cells showed complete synapsis of the autosomes, and 41% contained autosomes with discontinuous or no synapsis, indicating that HEIP1 is required for complete autosomal synapsis (Figure 2F). Remarkably, the presence of extended synapsis observed in *Heip1^-/-^*mutant is compatible with the residual loading of MSH4 observed in the mutant (see Figure 2C for residual loading of MSH4 in *Heip1^-/-^* mutant). Moreover, in 24% of spermatocytes, we observed synapsis between multiple chromosomes, suggesting that HEIP1 may prevent non-homologous synapsis (Figure 2E). Synapsis of the XY chromosomes was severely defective, with unpaired chromosomes being observed in 80% of pachytene cells (Figure 2E and G, arrowheads show unsynapsed sexual chromosomes).

### Dynamic chromosomal localization of HEIP1 during meiotic prophase I

To better understand how HEIP1 regulates other pro-CO proteins, we investigated its localization in wild type spermatocytes during meiotic prophase I. HEIP1 appeared first at early zygonema as punctate foci on synapsed axes (Figure 3A). We also observed few discrete foci on the axial elements of unsynapsed axes (Figure S3A, arrowheads). Numbers of HEIP1 foci increased during zygonema as synapsis progressed, peaking at late zygonema (120±20 foci) to early pachynema (103±16 foci) (Figure 3A). At these stages, HEIP1 appeared as punctate foci along the entire length of the synapsed axes. Super-resolution imaging of HEIP1 and SYCP3 by stimulated emission depletion (STED) microscopy resolved the two axes of synapsing chromosomes at zygonema and revealed that HEIP1 foci localized between the homologous axes even before they attain tight synapsis at a distance of 100 nm (Figure 3B, arrowheads). At early pachynema, HEIP1 foci were predominantly localized in the SC central region (Figure 3B). Lastly, numbers of HEIP1 foci decreased as pachynema progressed and only one or two foci per chromosome remained at mid/late pachynema (Figure 3A), equivalent to the number of CO sites detected by MLH1 immunostaining (Figure 1E). At diplonema, HEIP1 staining had completely disappeared.

**Figure 3.**
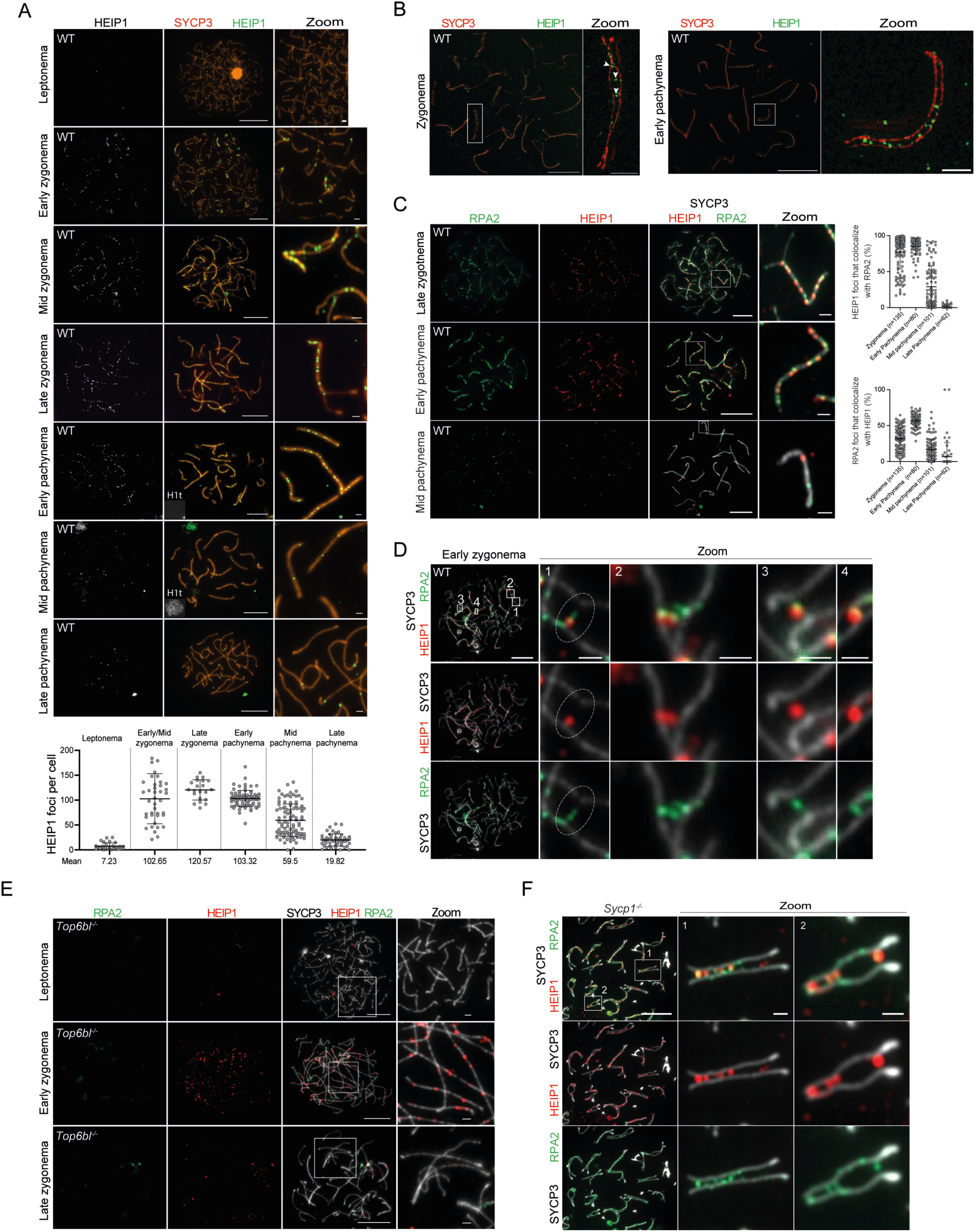
HEIP1 localization dynamics in prophase I spermatocytes. (A) HEIP1 localization in wild type spermatocyte nuclei at the different prophase I substages. Top: Representative images of chromosome spreads from prophase I spermatocyte nuclei immunostained for SYCP3 (orange), HEIP1 (green) and histone H1t (grey). Bottom: quantification of HEIP1 foci per spermatocyte nucleus at the different prophase I stages. The mean numbers of foci are shown below. Numbers of nuclei analyzed: 30, 40, 21, 77, 29, 96 and 52 for leptonema, early/mid zygonema, late zygonema, early pachynema, mid pachynema, late pachynema, respectively. (B) HEIP1 localizes to the central region of the synaptonemal complex. STED microscopy images of chromosome spreads from wild type spermatocytes at late zygonema and early pachynema immunostained for SYCP3 (red) and HEIP1 (green). Arrowheads highlight HEIP1 foci between chromosome axes. White squares correspond to the zoom images that show representative chromosomes. (C) HEIP1 colocalizes with RPA2 in prophase I spermatocytes. Left: Representative images of prophase I chromosomes from wild type mice immunostained for SYCP3 (grey), HEIP1 (red) and RPA2 (green). Right: Quantification of HEIP1 colocalization with RPA2 (top) and RPA2 colocalization with HEIP1 (bottom). (D) Localization dynamics of HEIP1 and RPA2 in wild type spermatocytes at early zygonema. Four steps of HEIP1 and RPA2 dynamics during synapsis are shown in the enlarged images 1 to 4 from the white squares. (E) Chromosome spreads from *Top6bl^-/-^*prophase I spermatocyte nuclei immunostained for SYCP3 (grey), HEIP1 (red) and RPA2 (green). (F) Chromosome spreads from 60 dpp *Sycp1^-/-^* prophase I spermatocyte nuclei immunostained for SYCP3 (grey), HEIP1 (red) and RPA2 (green). For each panel, magnified images show representative chromosomes. Scale bars, 10 μm for full nuclei and 1 μm for magnified images (2 µm for (B)). On graphs, bars indicate the mean ± SD and n=number of nuclei. Unless stated, 18 dpp mice were used.

### HEIP1 co-localizes with the ssDNA protein RPA2 but not with the recombinase DMC1

To test whether the HEIP1 localizes to recombination sites, spermatocyte chromosome spreads were costained for HEIP1 and RPA2 (Figure 3C and Figure S3A-B). Strikingly, at zygonema and early pachynema, 77% and 85% of HEIP1 foci, respectively, colocalized with RPA2 foci, indicating that most HEIP1 foci localize to HR intermediates that contain ssDNA (Figure 3C). Reciprocally, only a subset of RPA2 foci colocalized with HEIP1 (32% at zygonema and 57% at early pachynema), suggesting that HEIP1 is not present at all RPA2-coated ssDNA recombination intermediates (Figure 3C). In subsequent stages, numbers of HEIP1 and RPA2 foci decreased, and HEIP1-RPA2 colocalization decreased to just 30% at mid pachynema and 1% at late pachynema, implying that late intermediates marked by HEIP1 foci contain much less ssDNA (Figure 3C), as expected for intermediates progressing from single-end invasions to double Holliday junctions.

Colocalizing HEIP1-RPA2 structures were highly dynamic during early zygonema (Figure 3D and S3A). In unsynapsed regions, HEIP1-RPA2 structures localized close to an axis (Figure S3A, arrowheads on zooms 1, 2 and 3) or as nascent bridges that extended from an axis (Figure 3D; Zoom 1). As aligned axes progressed towards synapsis, HEIP1 localized with RPA2 bridges that moved from axis sites (Figure 3D; Zoom 2) towards the SC central region (Figure 3D; Zooms 3 and 4). These data suggest that HEIP1 follows RPA2/ssDNA-associated recombination intermediates as they bridge the axes of homologous chromosomes and mediates their synapsis.

To further define the specific recombination intermediates associated with HEIP1, colocalization with DMC1 was examined (Figure S3C-E). DMC1 foci appeared earlier than HEIP1, as shown by the presence of multiple DMC1 foci on chromosome axes during leptotene (Figure S3C). At zygonema, when HEIP1 first appeared, only 17% of foci colocalized with DMC1 foci (Figure S3E). However, among the HEIP1-DMC1 foci, a small fraction is located at the synaptic forks and on synapsed regions. These localization dynamics may reflect a transient overlap between DMC1 and HEIP1 as ssDNA is formed at nascent D-loops (Figure S3E, arrowheads).

### Initial localization of HEIP1 is DSB independent

In wild-type spermatocytes, HEIP1 strongly colocalized with RPA2 already at zygonema, suggesting a connection between HEIP1 recruitment and early steps of recombination. To further investigate this relationship, we monitored whether HEIP1 recruitment depends on DSB formation. *Top6bl^-/-^* mice lack the B-subunit of the SPO11- TOPOVIBL core DSB complex and are completely defective for DSB formation but can still form extensive synapsis although this occurs between non-homologous chromosomes (5) (Figure 3E, Figure S3F). In *Top6bl^-/-^* spermatocytes with well-developed lateral elements but no non-homologous synapsis, there were on average 90±37 HEIP1 foci, distributed along the entire lengths of the unsynapsed chromosome axes. Importantly, we did not detect any RPA2 foci in *Top6bl^-/-^* spermatocytes, as expected if breaks are not formed (Figure 3E, Figure S3F). HEIP1 loading on the axis can therefore occur independently of DSB formation. In later stage in *Top6bl^-/-^* spermatocytes, when partial non-homologous synapsis has occurred, numbers of HEIP1 foci were globally diminished suggesting that a DSB-independent pathway removes HEIP1 from the chromosome axes (Figure 3E, Figure S3F).

SPO11-TOPOVIBL-catalyzed DSB formation requires several axis-associated accessory factors including the REC114-MEI4-IHO1 complex and MEI1 (3, 51). As recruitment of these factors occurs independently of DSB formation, we wondered whether they play a role in HEIP1 localization. To this end, HEIP1 localization was analyzed in *Rec114^-/-^*spermatocytes (Figure S3G). The mean number of HEIP1 foci on chromosome axes was 123±54, revealing that HEIP1 loading can also occur in the absence of REC114 (Figure S3G).

### Synapsis is not required for repositioning of HEIP1 structures

HEIP1-RPA structures relocalize from axis-associated positions to the central region of the SC as synapsis ensues (Figure 3D). To determine whether the SC central element is required for HEIP1 to reposition, we monitored HEIP1 and RPA2 localization in *Sycp1^-/-^* mice that fail to synapse due to absence of the SC central-region transverse filament protein SYCP1 (Figure 3F) (41). In *Sycp1^-/-^* spermatocytes, homologs align but fail to synapse at a distance of 100 nm. While meiotic recombination initiates normally, progression is delayed and CO sites fail to mature, indicated by the absence of MLH1 foci (41). HEIP1 foci were detected in *Sycp1^-/-^*zygotene-like spermatocytes with well-developed axes and many foci were located between homolog axes and colocalized with RPA2 (Figure 3F). This finding indicates that synapsis is not required for (i) HEIP1-RPA2 structures to reposition from on-axis to between-axes positions, and (ii) HEIP1 to colocalize with RPA2-associated recombination intermediates.

### HEIP1 colocalizes with crossover factors TEX11 and MLH1

To further understand the link between HEIP1 and class I COs, we investigated HEIP1 colocalization with known class I CO factors (TEX11 and MLH1). TEX11 is a TPR repeat protein and ortholog of budding yeast Zip4 that acts as a platform for other CO proteins and facilitates CO formation by stabilizing HR intermediates (7). In wild-type spermatocytes, TEX11 formed immunostaining foci along synaptonemal complexes during zygonema and pachynema (Figure 4A). At early pachynema, 81% of TEX11 foci colocalized with HEIP1 foci (Figure 4A, B). Reciprocally, only 55% of HEIP1 foci colocalized with TEX11 at this stage, i.e. HEIP1 foci are in excess relative to TEX11 possibly reflecting earlier loss of TEX11 from chromosomes (Figure 4A, B). The persistent subset of HEIP1 foci at mid-pachynema is temporally and numerically equivalent to that of CO maturation factors, such as the MutL*γ* proteins MLH1 and MLH3. Indeed, 80% of MLH1 foci co-localized with HEIP1, suggesting a role for HEIP1 in CO maturation (Figure 4C, D). Notably, not all HEIP1 foci gave rise to COs because only 57% are colocalized with MLH1 (Figure 4C, D). At late pachynema, 49% of MLH1 foci colocalized with HEIP1 (Figure 4D) reflecting disappearance of HEIP1 at this substage and suggesting that its role is completed before the crossing over is implemented.

**Figure 4:**
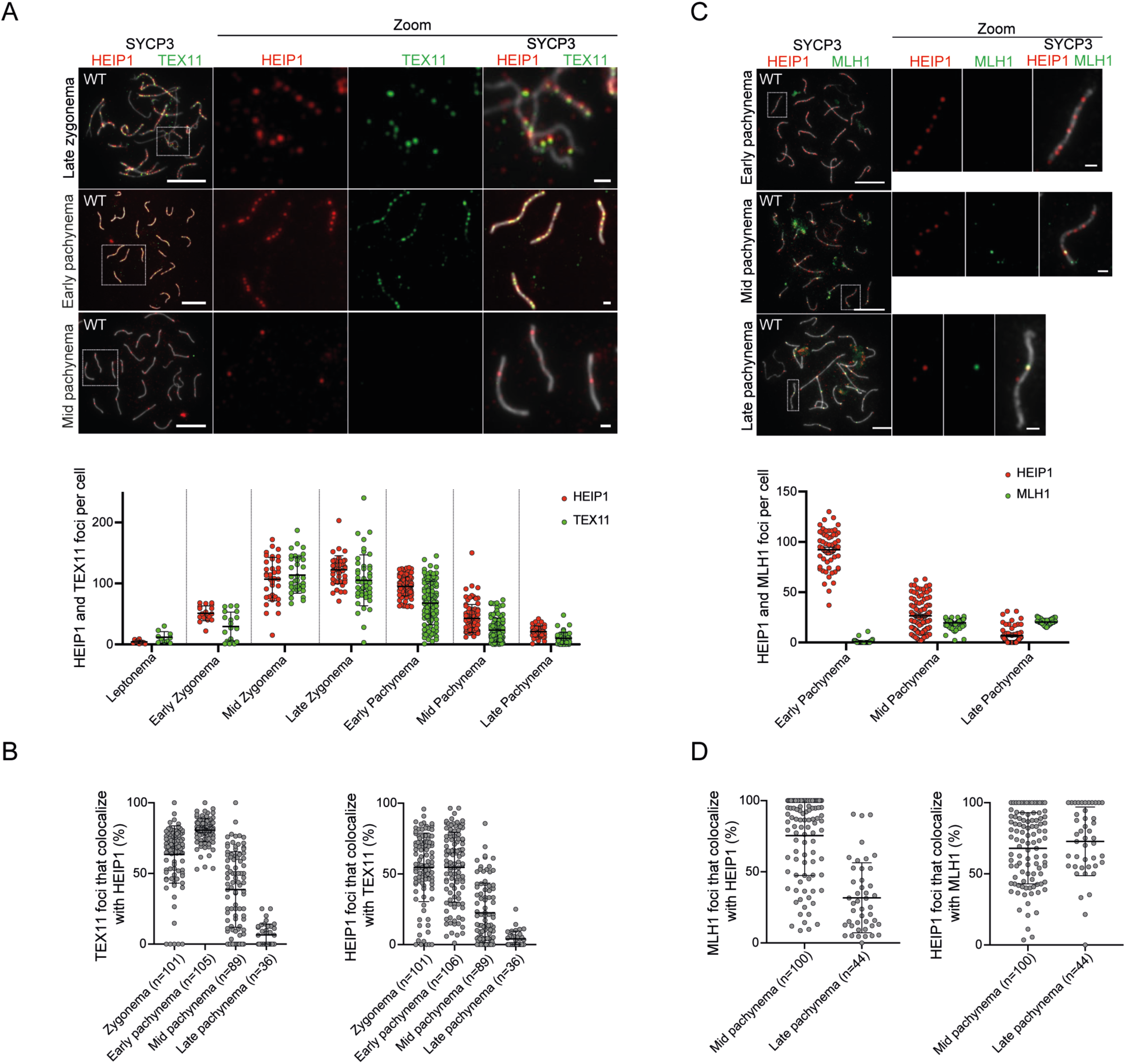
HEIP1 progressively localizes to crossover sites. (A) HEIP1 colocalization with TEX11 at zygonema and pachynema. Upper panels: immunostaining of wild type spermatocytes for SYCP3 (grey), HEIP1 (red) and TEX11 (green). on the right. Bottom: quantification of HEIP1 and TEX11 foci during prophase I. (B) Quantification of TEX11 colocalization with HEIP1 and *vice versa*. (C) HEIP1 colocalization with MLH1 at pachytene stages. Upper panels: immunostaining for SYCP3 (grey), HEIP1 (red) and MLH1 (green) of wild type pachytene spermatocyte nuclei. Bottom: quantification of HEIP1 and MLH1 foci during prophase I. (D) Quantification of MLH1 colocalization with HEIP1 and *vice versa*. 18 dpp mice were used; on each panel magnified images (zoom) of representative chromosomes (white boxes) are presented, Scale bars, 10 μm for full nuclei and 1 μm for magnified images. On graphs, bars indicate the mean ± SD.

### HEIP1 physically interacts with the E3 ligase HEI10

Our results suggest a conserved role for HEIP1 in mammals, similar to that described for plant HEIP1. To test whether mammalian HEIP1 also interacts with E3 ligase HEI10, as shown in plants (31, 57), co-immunoprecipitation (co-IP) and yeast two-hybrid experiments were performed (Figure 5A and Figure S4A). Of note, we thus confirmed that HEIP1 protein is undetectable on protein extract from *Heip1^-/-^* testes. Both assays detected a clear interaction between HEIP1 and HEI10. The yeast two-hybrid assay showed that interaction occurs between a HEIP1 N-terminal fragment (1–226) and the HEI10 C-terminal region (81–276) (Figure S4A).

**Figure 5.**
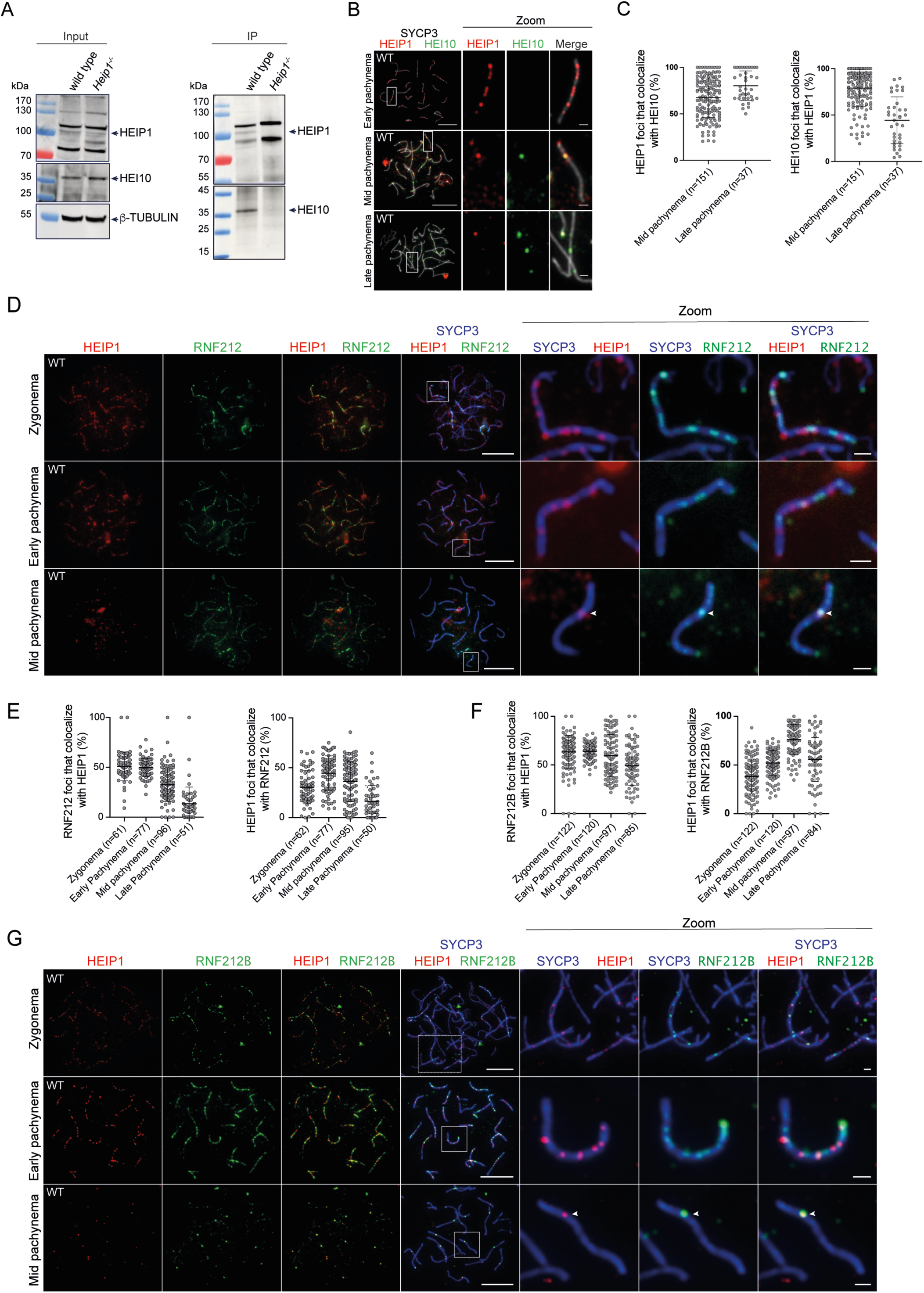
Interplay between HEIP1 and the pro-CO RING finger E3 ligases HEI10, RNF212 and RNF212B. (A) Co-immunoprecipitation of HEIP1 and HEI10 using wild type and *Heip1^−/^*^−^ testis protein extracts. Western blot inputs (left) and immunoprecipitation (right; IP) with rabbit antibodies against the C-terminal part of HEIP1. HEIP1 band in the wild type is detected at a size corresponding to a molecular weight of 100 KDa instead of 73 KDa expected, which could be the consequence of (i) potential posttranslational modifications (ii) or of the intrinsic biochemical property of the protein. β-tubulin was used as input control. (B) HEIP1 colocalization with HEI10 at pachytene stages. Immunostaining for SYCP3 (grey), HEIP1 (red) and HEI10 (green) in wild type spermatocytes at early, mid and late pachynema. (C) Percentage of HEIP1 foci that colocalize with HEI10 foci, and *vice versa*. (D) HEIP1 colocalization with RNF212. Immunostaining for SYCP3 (blue), HEIP1 (red) and RNF212 (green) on wild type spermatocytes at zygonema, early and mid-pachynema. Arrowheads show CO sites. (E) Percentage of RNF212 that colocalizes with HEIP1, and *vice versa*. (F) Percentage of RNF212B foci that colocalize with HEIP1 foci, and *vice versa*. (G) HEIP1 colocalization with RNF212B. Immunostaining for SYCP3 (blue), HEIP1 (red) and RNF212B (green) on wild type spermatocytes at zygonema, early and mid-pachynema. Arrowheads show CO sites. For all images: zoom are magnified images of representative chromosomes (white boxes); scale bars: 10 μm for full nuclei and 1 μm for magnified images. For each graph: bars indicate the mean ± SD. 18 dpp mice were used.

### HEIP1 colocalizes closely with HEI10 and partially with RNF212 and RNF212B

To further explore the relationship between HEIP1 and HEI10, we investigated their colocalization (Figure 5B, C and Figure S4B). 67% and 80% of HEIP1 foci colocalized with HEI10 respectively at mid and late pachynema. Conversely, 79% of HEI10 foci colocalized with HEIP1 foci at the mid-pachytene stage. But, as observed with MLH1, HEIP1 foci disappeared earlier than HEI10 foci, with the percentage of HEI10 that colocalized with HEIP1 decreasing to 44% at late pachynema (Figure 5C), suggesting that HEIP1 is not required for the last steps of CO maturation.

We also compared the colocalization of HEIP1 with the two other CORs, RNF212 and RNF212B (Figure 5D-G). HEIP1 and RNF212 appeared at zygonema and started to disappear during pachynema (39) (Figure 5D, E and Figure S4C). Only 45% of HEIP1 foci colocalized with RNF212 (and less than 50% of RNF212 foci colocalized with HEIP1) when HEIP1 foci reached their maximum number in early pachynema (Figure 5E). In addition, some late HEIP1 foci, which are expected to mark COs sites at mid-pachynema, colocalized with RNF212 foci (Figure 5D, arrowheads on the zoomed images), suggesting a cooperation between these two proteins at later stages of CO formation.

Previous studies showed that RNF212 and RNF212B overlap but do not show perfect colocalization, suggesting that these two proteins may have distinct functions during CO formation (33, 34). Therefore, we compared HEIP1 and RNF212B localization during prophase I of meiosis (Figure 5F, G). As seen for RNF212, HEIP1 showed only a partial colocalization with RNF212B at zygonema (38% of HEIP1 foci) and early pachynema (52% of HEIP1) (Figure 5F), suggesting that HEIP1 and RNF212B may have distinct roles during the early stages of meiotic recombination. At mid-pachynema, 76% of HEIP1 foci colocalized with RNF212B foci (Figure 5F, G, arrowheads on the zoomed images), suggesting that HEIP1 and RNF212B act collectively during MutLg- dependent steps of crossing over. At late pachynema, HEIP1 started to disappear, whereas RNF212B was still present at CO sites (Figure 5F, G), suggesting that HEIP1 and RNF212B do not collaborate for the last steps of CO formation. Remarkably, we observed a similar localization pattern respective to the different substages of prophase I for RNF212 and HEIP1.

Like HEIP1, RNF212 and RNF212B also are loaded on chromosomes independently of DSB formation but, their localization is restricted to the non-homologous synapsed regions (33, 34, 39). Moreover, it was proposed that DSB formation facilitates RNF212B recruitment on the SC but not RNF212 (34). We then compared the patterns of HEIP1 and RNF212 on meiotic chromosomes in the absence of DSBs (Figure S4E). Unlike HEIP1, which was concentrated along non-synapsed axial elements in early zygotene-like stage nuclei (Figure 3E and Figure S4E), RNF212 was restricted to non-homologously synapsed regions in late zygotene-like nuclei, as previously reported (34). Therefore, in the absence of DSBs, only 7% and 19% of HEIP1 foci colocalized with RNF212 in early and late zygonema-like nuclei, respectively (Figure S4E). Knowing that RNF212B also colocalize with non-homologous synapsed regions, with a partial DSB formation dependency(34), we envision that in the *Top6bl^-/-^* mutant the spatiotemporal deposition patterns of RNF212/RNF212B and HEIP1 are mediated by distinct mechanisms.

### HEIP1 orchestrates the recruitment of the E3 ligases HEI10, RNF212 and RNF212B on meiotic chromosomes

The identification of HEI10 as an interacting partner of HEIP1 in mammals, and the spatial and temporal correlations between HEIP1, HEI10, RNF212B and to a lesser extent RNF212, suggest a functional relationship between HEIP1 and the COR proteins.

Consistently, formation of crossover-specific HEI10 foci was promoted by HEIP1 (Figure 6A). The mean number of HEI10 foci at mid-pachynema was reduced from 22±3 in wild-type spermatocytes to 4.4±4 in *Heip1^-/-^* spermatocytes (Figure 6A). Dynamics of RNF212 and RNF212B foci were also facilitated by HEIP1. As previously described (26, 33, 34, 39), in wild type spermatocytes, RNF212 and RNF212B first appeared as numerous foci on synapsed chromosomes, reaching a maximum of ∼143±28 and 101±24 foci, respectively, at early pachytene (Figure 6B). RNF212 foci were then progressively less numerous, remaining at CO sites (Figure 6B) and disappearing completely at diplonema. The reduction of RNF212 and RNF212B foci to produce a crossover-specific pattern has previously shown to be HEI10 dependent (26). Similarly, in *Heip1^-/-^*spermatocytes, higher numbers of RNF212 and RNF212B foci persisted throughout pachynema, with respective means of 138±34 RNF212 and 98±32 RNF212B foci in mid pachynema (*versus* 69±39 and 49±28 in wild type cells) (Figure 6B). Focus numbers were even in late pachynema *Heip1^-/-^* spermatocytes with means of 110±37 RNF212 and 83±29 RNF212B foci (*versus* 36±16 for both proteins in wild-type). Focus numbers then started to decrease after pachynema but some RNF212 and RNF212B foci persisted until diplonema in *Heip1^-/-^* spermatocytes, whereas they completely disappeared in wild type (Figure S4F), indicating that normal dissociation of RNF212 and RNF212B from the chromosomes depends on HEIP1.

**Figure 6.**
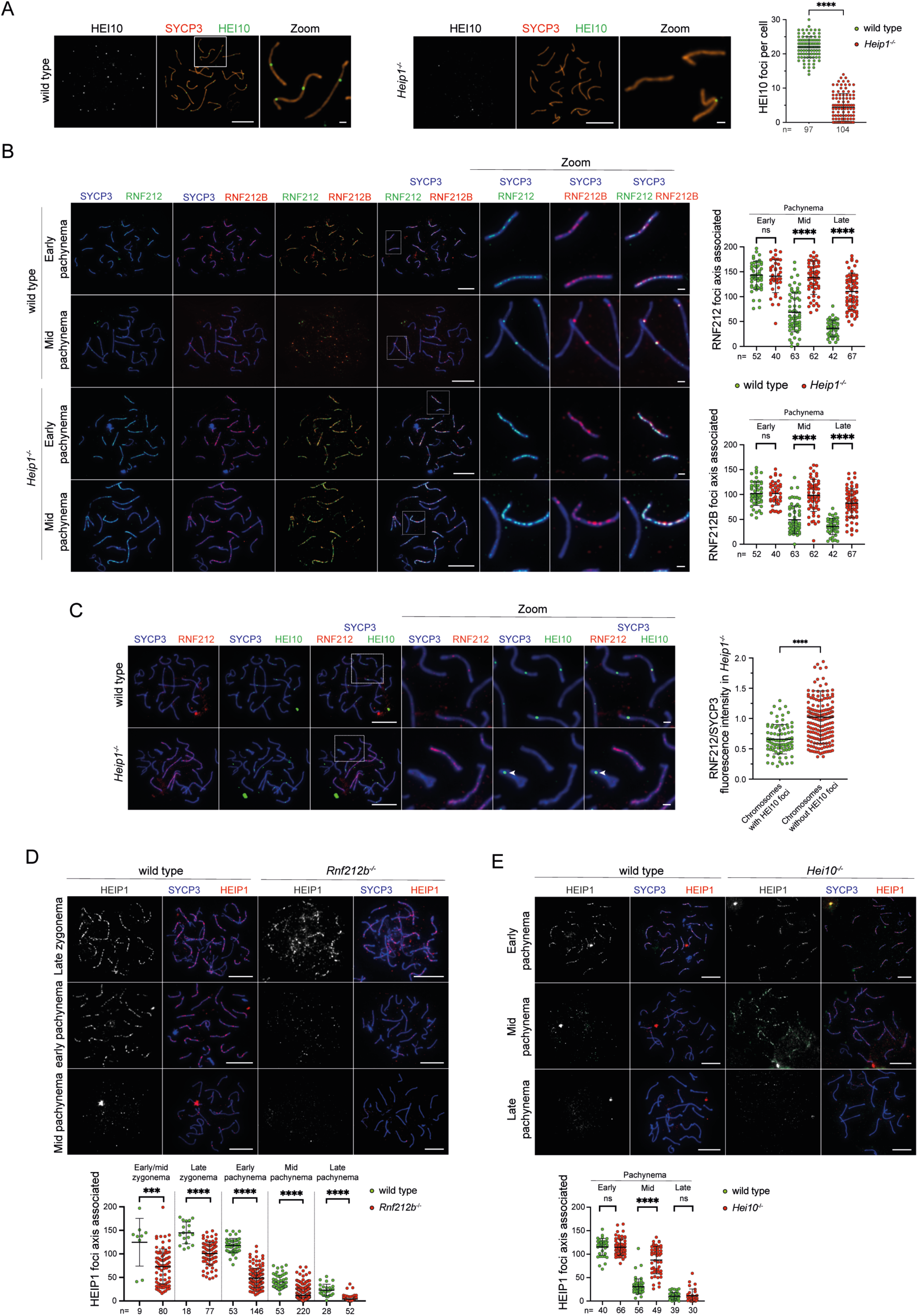
**Interdependent relationship between HEIP1 and the pro-CO RING finger E3 ligases.** (A) Chromosomal localization (left and middle) and quantification (right) of HEI10 in wild type and *Heip1^-/-^* mice. Immunostaining for SYCP3 (orange) and HEI10 (green) on wild type and *Heip1^-/-^* spermatocytes at mid pachynema. (B) Chromosomal localization (left) and quantification (right) of RNF212 and RNF212B in *Heip1^-/-^* mice. Immunostaining for SYCP3 (blue), RNF212 (green) and RNF212B (red) on wild type and *Heip1^-/-^* spermatocytes at early and mid-pachynema. (C) Left: chromosomal localization of RNF212 and HEI10 in *Heip1^-/-^* mice. Immunostaining for SYCP3 (blue), HEI10 (green) and RNF212 (red) on wild type and *Heip1^-/-^* spermatocytes at mid-pachynema. Right: ratio of the fluorescence intensity of RNF212 and SYCP3 on chromosomes with at least one HEI10 focus and on chromosomes without HEI10 foci. (D) Chromosomal localization (top) and quantification of HEIP1 (bottom) in wild type and *Rnf212b^-/-^* mice. Immunostaining for SYCP3 (blue) and HEIP1 (red) on wild type and adult *Rnf212b^-/-^* spermatocytes at late zygonema, early and mid- pachynema. (E) Chromosomal localization (top) and quantification (bottom) of HEIP1 in *Hei10^-/-^*mice. Immunostaining for SYCP3 (blue) and HEIP1 (Red) on adult wild type and *Hei10^-/-^* spermatocytes at early, mid and late pachynema. 18 dpp mice were used, otherwise stated. Zoom: higher magnification of representative chromosomes (white boxes). Scale bars, 10 μm for full nuclei and 1 μm for magnified images. Bars indicate the mean ± SD; ***P < 0.001, ****P < 0.0001 (two- tailed unpaired Mann-Whitney test) and on each graph *n*= numbers of analyzed nuclei.

We detected residual HEI10 foci in a few *Heip1^-/-^* nuclei (Figure 6C, arrowheads) and observed that HEI10-positive chromosomes had 1.6-fold lower RNF212 fluorescence intensity than HEI10- negative chromosomes. Moreover, qualitative analysis revealed that on the HEI10-negative chromosomes, the RNF212 signal was diffuse along the SC, while HEI10-positive chromosomes accumulated RNF212 as foci that colocalized with HEI10 (Figure 6C). Collectively, these data suggest that HEIP1 regulates RNF212 and RNF212B turn-over by directly controlling HEI10 recruitment in *cis*, and as such may acts as a master regulator of COR dynamics.

Finally, we determined whether the CORs reciprocally influence HEIP1 localization. Numbers of HEIP1 foci were significantly reduced in *Rnf212b^−/−^*mutant spermatocytes at early pachynema (41% of wild-type), despite being present on axes at earlier stages (Figure 6D) suggesting that RNF212B (and possibly RNF212) is important to stabilize HEIP1 association. Conversely, in *Hei10^−/−^* spermatocytes, HEIP1 accumulated as pachytene progressed, with 2.8-fold more foci that wild type by mid pachynema, indicating that HEI10 promotes the turnover of HEIP1 (Figure 6E). Together, these observations point to crosstalk between the CORs and HEIP1.

## Discussion

### HEIP1 is an evolutionarily conserved pro-CO factor

The identification and characterization of meiotic pro-class I CO factors remain an essential goal to fully understand the regulation of this activity that is essential for gametes formation, genetic stability and diversity. Here, we provide an extensive characterization of HEIP1, a conserved pro-class I CO factor, in the mouse. We found that HEIP1 acts as a master regulator of CO formation by interacting with the meiotic chromosomes throughout prophase I, first independently of HR initiation and then by following recombination intermediates independently of SC central element formation. At the molecular level, we demonstrated that HEIP1 directly interacts with HEI10 and orchestrates the proper spatiotemporal recruitment of different pro-CO factors during prophase I. We thus propose that it acts as a recruitment platform protein for different recombination factors.

From our data, we propose that HEIP1 is a conserved class I pro-CO factor in the mouse. First, *Heip1^-/-^* mice are sterile and show a strong decrease of CO formation, although DSB formation and the initial HR steps are not affected. Second, the loading of the pro-CO factors MSH4 and TEX11 is significantly reduced, suggesting that recombination intermediates are less prone to mature into COs due to reduced MutS*γ*-dependent stabilization. Even if we could not detect any significant difference in Bloom recruitment in the absence of HEIP1, we propose that in the absence of HEIP1, nascent unprotected recombination intermediates are dissociated by the Bloom complex, which is presumed to promote NCO formation by synthesis-dependent strand annealing (11, 19, 47, 48) . Third, HEIP1 exhibits a spatiotemporal localization compatible with a class I pro-CO factor, similar to what described in rice. Initially, HEIP1 appears as numerous foci during the early stages of prophase I (leptonema in rice, early zygonema in the mouse) and gradually becomes localized to MLH1/HEI10-dependent class I CO sites at pachynema (31). We also found that these foci perfectly colocalize with the pro-CO proteins TEX11, MLH1 and HEI10, suggesting that it assists DNA intermediates until CO formation. However, HEIP1 does not persist as long as MLH1 or HEI10, suggesting that it is not required for the last steps of CO formation. Lastly, several data, including from this study, demonstrate that HEIP1 binds to the same partners in plants and mammals. Yeast two-hybrid assays showed that rice HEIP1 interacts with other pro-CO proteins, such as OsHEI10, OsZIP4, and OsMSH5, although this finding needs to be confirmed *in vivo* (31, 58). Here, we found that HEIP1 interacts with HEI10 *in vivo* and by yeast two-hybrid assay. Moreover, both proteins co- localize at the pachytene stage and HEI10 loading requires HEIP1. These results indicate that HEIP1 directly collaborates with HEI10 to promote CO formation. In line with our data, it was proposed that REDIC1/HEIP1 interacts with MSH5 and ZIP4/TEX11 *in vitro* (37). These results suggest that despite its highly divergent amino acid sequence, mouse HEIP1 also interacts with other pro-CO proteins. HEIP1 sequence was identified in various plants and animals and the conservation of the interactions between HEIP1 and pro-CO proteins suggests that they may be present in the last common ancestor of eukaryotes species harboring the *HEI10*, *MSH5* and *TEX11/ZIP4* genes.

### Bimodal regulation of HEIP1 dynamics during prophase I of meiosis

We propose a two-step regulation of HEIP1 spatiotemporal localization. The first step is defined by HEIP1 loading as discrete foci on chromosome axes at early zygonema, before or at the time of DSB formation and HR initiation and largely before CO resolution. This early HEIP1 recruitment is also observed in rice where HEIP1 foci are detectable before synapsis formation, suggesting that the early HEIP1 deposition on chromosomes is evolutionarily conserved (31). Importantly, in *Top6bl^-/-^* and *Rec114^-/-^*mice, in which DSBs are not formed and homologous synapsis abolished, we observed HEIP1 loading on the axis, as discrete foci, specifically at the early zygotene-like stage. Remarkably, the specific timing of recruitment of HEIP1 in DSB formation mutants that we observe could explain the discrepancy with the previous characterization of HEIP1 (REDIC1) recruitment performed by Fan *et al.* in the *Top6bl^-/-^*mutant. Indeed, in their study the authors did not detect HEIP1 in the mutant, but they focused on late zygotene stages at which the protein is already removed (see Figure 3E and S3G). Altogether, this result has two implications. First, since in these mutants, unsynapsed axes persist longer than in wild type, a transient interaction of HEIP1 with the chromosome axes may be more likely to detect than in wild type spermatocytes, although such foci can be seen on wild type unsynapsed chromosomes (Figure S3A). We then believe that the TOPOVIBL or REC114 depletion triggers the accumulation of HEIP1 on the chromosome axis. Second, at the physiological level, it suggests that meiotic recombination initiation is not required for the initial HEIP1 loading.

One unanswered key question is how HEIP1 interacts with the axis. One possibility is that HEIP1 recruitment is triggered by axial element components, the loading of which is independent of DSB formation (*e.g.* SYCP3, HORMAD1/2) (59). Our results showed that HEIP1 loading is independent of the core DSB catalytic complex (TOPOVIBL/SPO11) and of the DSB accessory factor REC114, which is loaded on the axes in a DSB formation-independent manner. REC114 is part of the RMM complex (REC114-MEI4-IHO1). REC114 and MEI4 form a subcomplex, their absence leads to similar phenotypes and their localization is interdependent, but independent of IHO1. IHO1, which interacts with HORMAD1, may act as an axis anchor protein that triggers the recruitment of partners. Therefore, it would be interesting to test if the initial HEIP1 loading relies on IHO1. Interestingly, post-translational modifications are present on the axes, at least in the yeast and mouse (60–62), and they might also participate in regulating the first step of HEIP1 targeting.

HEIP1 is not the only pro-CO factor associating with chromosomes independently of DSB formation. RNF212 and, to a lesser extent, RNF212B also are present on *Spo11^-/-^* chromosomes (33, 34, 39). However, as opposed to HEIP1, the E3 ligases RNF212 and RNF212B specifically localize on synapsed regions between non-homologous chromosomes, suggesting that HEIP1 and the E3 ligases RNF212 and RNF212B have distinct DSB-independent mode of recruitment and regulation. This leads to two non-exclusive hypotheses. The first is that HEIP1 and RNF212/RNF212B exhibit an HR-independent pattern of regulation, but with a different pattern of loading on the chromosome, which reflects the requirement of the early recruitment of the proteins that designate class I CO sites. The second hypothesis is that these proteins have an early role during prophase I, independent of class I CO designation and resolution.

Following the initial HR-independent recruitment of HEIP1, we propose that the second step of regulation of HEIP1 spatiotemporal localization, which is HR-dependent, is initiated by its recognition and interaction with recombination intermediates. First, we found that in the absence of TOPOVIBL and REC114, HEIP1 discrete foci are removed at the late zygotene-like stage when synapsis between non-homologous chromosomes starts. This suggests that HEIP1 maintenance on prophase I chromosomes, and particularly on the SC, relies on DSB formation and HR. Second, in wild type spermatocytes, HEIP1 axis-associated foci co-localize with RPA2 foci, suggesting a relationship between HEIP1 and RPA2-coated ssDNA structure. To explain its stabilization after DSB formation, we propose that HEIP1 binds directly to ssDNA, or that it interacts with proteins of the recombination machinery. Supporting our hypothesis, a direct *in vitro* interaction between REDIC1/HEIP1 with branched DNA structure was previously reported (37). Surprisingly, HEIP1 foci co-localize poorly (20% at zygotene stage) with DMC1 (strand exchange protein) foci. Thus, the high degree of co-localization with RPA2 and not with DMC1 leads us to propose that HEIP1 is interacting with recombination intermediates prior to DMC1/RAD51 coating and strand exchange reaction (Figure 7A (a)) or later by binding to single strand DNA at or before the second-end DNA capture (Figure 7A (b)). In both cases, we propose that HEIP1 location might help the second-end DNA capture formation promoted by the combined activity of RAD52 and RPA2 (63) and the recruitment/orchestration of pro-CO factors, such as the MutS*γ* and ZZS complexes (7, 45, 64, 65). However, future experiments will be required to rigorously identified the precise DNA/protein structure recognized by HEIP1. Finally, we propose that this progressive HEIP1-dependent stabilization of DNA intermediates leads to the formation of dHJs that will be processed as COs by the MutL*γ* complex.

**Figure 7.**
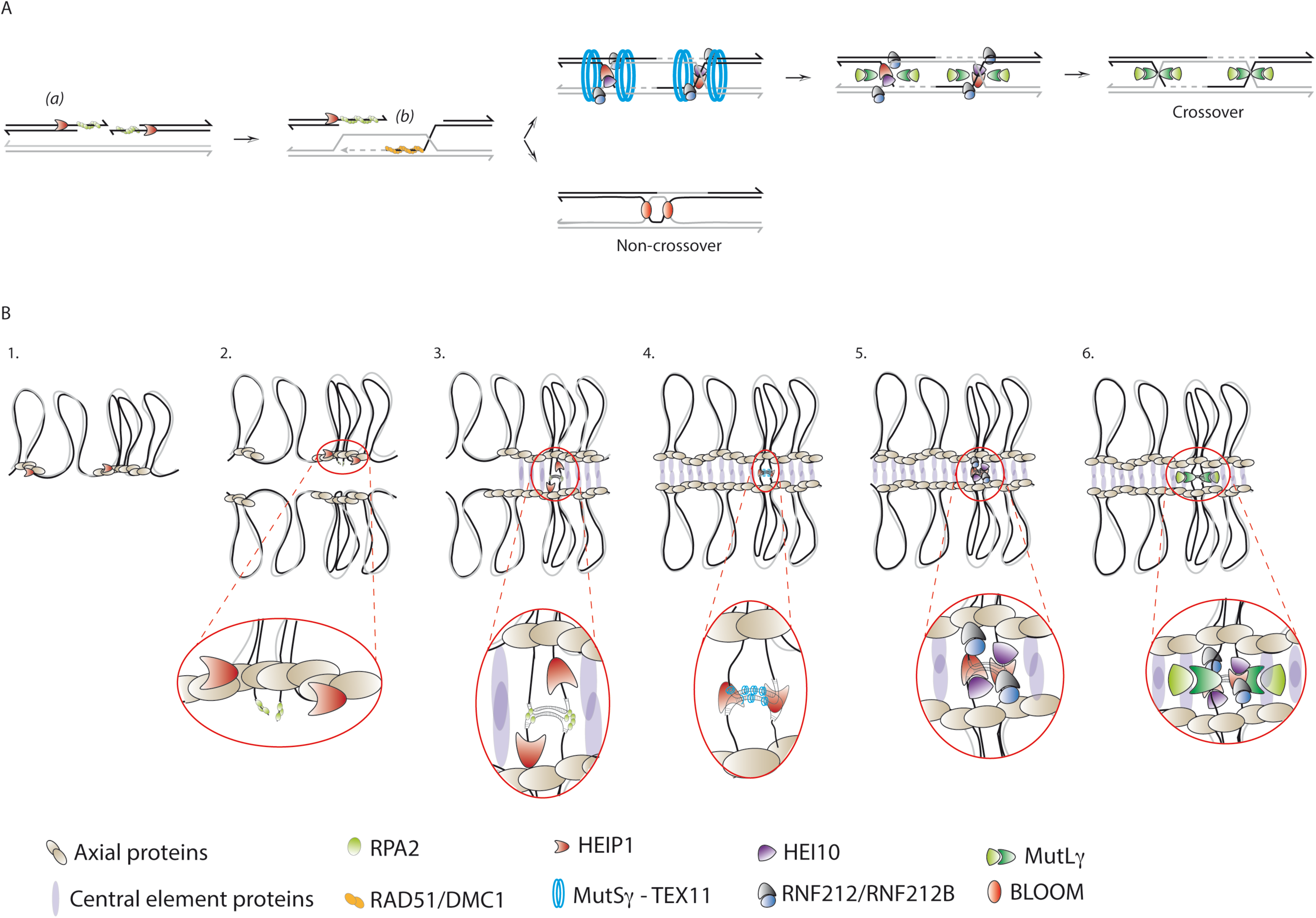
Model for HEIP1-dependent CO orchestration during prophase I of meiosis. Proposed HEIP1 roles during homologous recombination (A) and model that illustrates HEIP1 localization on chromosomes (B). (A) After break formation, HEIP1 co-localizes with RPA2 recombination intermediates, prior to DMC1/RAD51 (a) loading and/or before second end capture (b). At these positions, HEIP1 may promote second-end capture for repair and recruit CO- promoting factors, such as the MutSγ complex and TEX11 (blue circles), to protect recombination intermediates against Bloom dissolution (NCO). Simultaneously, HEIP1 facilitates RNF212 and RNF212B relocalization, HEI10 and MutL*γ* recruitment. Then, at the time of crossover resolution, HEIP1 is removed. (B) (1) HEIP1 (red shapes) initially appears along chromosome axes independently of DSBs. (2) During chromosome alignment, HEIP1 relocates and colocalizes with single-stranded DNA (ssDNA) coated by RPA2 (green circles), marking recombination sites. (3) During chromosome synapsis, HEIP1 transitions from the axis to the central region of the synaptonemal complex (SC). (4) At these positions, HEIP1 may recruit CO-promoting factors, such as the MutSγ complex and TEX11 (blue circles). (5–6) Then, HEIP1 presence promotes the proper deposition of HEI10, RNF212, RNF212B, and MutL*γ*, which will mark the future crossover sites. Once all proteins required for CO maturation are recruited, HEIP1 disappears.

### HEIP1 promotes CO formation by orchestrating the activity of different pro-CO proteins

Our study demonstrates that HEIP1 plays a pivotal role in CO maturation. First, HEIP1 strongly co- localizes with TEX11 and MSH4 (see also (37)), two proteins proposed to stabilize early recombination intermediates, such as D-loops. Moreover, the localization of TEX11 and MSH4 relies on HEIP1, suggesting that HEIP1 modulates the local recruitment of pro-CO proteins required for the early steps of CO formation. The HEIP1 structure is predicted to be largely disordered, suggesting that HEIP1 may behave like an intrinsically disordered protein, which relies on its intrinsic malleability to bind to and recruit partners (66, 67). Interestingly, TEX11 forms a complex along with SHOC1 and SPO16 (ZZS complex), while the MSH4-MSH5 heterodimer forms the MutS*γ* complex (46, 55, 65, 68). Biochemical studies suggest that both complexes act differently to stabilize early recombination intermediates (8, 10, 68). As HEIP1 is likely to interact with TEX11 and MSH5 (37), we propose that HEIP1 stabilizes early recombination intermediates by coordinating different biochemical properties carried by the ZZS and MutS*γ* complexes. Furthermore, we propose that this early HEIP1 function is independent of its interaction with HEI10.

In addition to this role in MutS*γ* and TEX11 stabilization, we uncover a later role of HEIP1 during CO formation, with the timing of HEIP1 localization coinciding with the last steps of CO maturation at late mid/late pachytene. Moreover, HEIP1 strongly colocalizes with the CO markers MLH1 and HEI10, and interacts physically with HEI10. This suggests that it could modulate the activity of enzymes involved during dHJ resolution as COs. HEIP1 role in recruiting HEI10 in plants and mice and the ability of disordered proteins to bind to several partners suggest that HEIP1 may influence the local concentration of HEI10 and other late CO factors at future CO sites, possibly by acting as a scaffold protein.

In *A. thaliana heip1* mutants, numbers of MLH1/HEI10 foci are decreased by only 20% but the number of chiasmata (the cytological view of COs) is reduced by 50% relative to wild type (32), suggesting an additional role of plant HEIP1 during CO maturation downstream of MLH1/HEI10 focus formation. Such a role in mice seems unlikely because HEIP1 foci at CO sites disappear before MLH1 and HEI10. Moreover, we observed a similar drastic reduction of chiasmata and of MLH1/HEI10 foci numbers in *Heip1^-/-^* spermatocytes, suggesting a divergence of function during CO maturation between organisms.

Altogether, these data suggest that HEIP1 is a master regulator of meiotic recombination that acts at several steps during CO formation, starting from the stabilization of early recombination intermediates until they form dHJs that will be resolved as CO.

In the mouse, the pro-CO RING finger E3 ligases RNF212, RNF212B and HEI10 are implicated in organizing meiotic recombination until CO formation. During CO formation, RNF212 and RNF212B are mutually dependent, and HEI10 loading requires both RNF212 and RNF212B. In turn, the timely removal of RNF212 and RNF212B relies on HEI10 (26, 33, 34). HEIP1 regulates the behavior of these proteins during meiotic prophase I. HEIP1 is important for RNF212 and RNF212B removal and for the loading of HEI10 on chromosomes. However, our data showed that *Heip1^-/-^* mutant does not phenocopy *Hei10^-/-^* mutant and, in particular, unlike what described in *Hei10^-/-^* mice by Qiao and colleagues (26), pro-CO proteins, such as MSH4, do not increase but rather decrease in the *Heip1^-/-^* mutant, suggesting that HEIP1 also acts earlier during CO formation by modulating (directly or indirectly) MSH4 pro-CO activity in a HEI10-independent manner. Therefore, HEIP1 does not only promote HEI10 recruitment.

We propose that HEIP1 also orchestrates the activity of the E3 ligases RNF212, RNF212B and HEI10 during CO formation. Recent studies revealed that RNF212B localization differs from that of RNF212. Specifically, the differentiation of CO-specific RNF212B foci occurs earlier and is significantly more pronounced compared with RNF212 foci (33, 34). In addition, RNF212 and RNF212B foci do not co-localize perfectly (60% of overlap). This is partly explained by the higher abundance of RNF212 compared with RNF212B (33, 34). Altogether, these data suggest that RNF212 and RNF212B may not exclusively form heterodimers, and that the progressive maturation of recombination intermediates into COs requires the dynamics of complexes involving switching partners between RNF212, RNF212B, and possibly HEI10. Interestingly, the colocalization of HEIP1 with HEI10, and partially with RNF212 and RNF212B, suggests that it could organize the switching of binding partners between E3 ligases and modulate post-translational modifications that are crucial to orchestrate the correct maturation of DNA intermediates (potentially by stabilizing MSH4-MSH5 heterodimers) until they are formed into mature COs.

### The HEIP1 acting model

We propose that HEIP1 is a caretaker of nascent recombination intermediates, until they are resolved as COs that act as a scaffold protein that spatio-temporally controls the recruitment different recombination factors. For this reason, we envision HEIP1 as a master regulator of crossover formation. First, at zygonema, HEIP1 is recruited on the chromosome axis through a yet-uncharacterized process, but seemingly independently of DNA DSB formation (Figure 7B (1)). Then, it relocates to ssDNA DSB ends coated by RPA2, which may represent recombination sites (Figure 7B (2)). During synapsis establishment, while DNA strand invasion occurs, HEIP1 relocates from the axis to the central region of the synaptonemal complex (Figure 7B (3)). Here, HEIP1 would exert its pro-CO activity by facilitating the recruitment of its partners, such as the MutS*γ* complex (through MSH5) and TEX11 (Figure 7B (4)), while recombination intermediates are already exposed to the Bloom/Sgs1-TOP3-RMI1 unwinding activity, as demonstrated in budding yeast (11, 20, 45, 69). This reinforcement of recombination intermediates by HEIP1 would also participate in consolidating chromosome synapsis between homologous chromosomes. Once homologous chromosomes are totally synapsed at pachytene, CO-prone DNA intermediates coated by HEIP1 and additional pro-CO factors (*e.g.* MutS*γ* and ZZS complexes) would be progressively selected among stable DNA molecules through the HEIP1- dependent recruitment of HEI10 and its partnership with RNF212 and RNF212B (Figure 7B (5)). The right combination of these E3 ligases and their activities would provide the adequate environment for DNA molecules to be processed as dHJs that will be further cleaved as COs by the MutL*γ* complex (Figure 7B (6)).

## Conclusion

Our study clearly identifies a conserved protein involved in CO formation, raising the total number of proteins required for class I CO formation to 17: TEX11, SHOC1, SPO16, M1AP1, HFM1, MSH4, MSH5, RNF212, RNF212B, HEI10, CDK2, CNTD1, PRR19, MLH1, MLH3, EXO1 and HEIP1. Recent models suggested that their concerted activities stabilize and shape recombination intermediates until CO formation. HEIP1, which is present at most steps of the HR pathway until CO formation and interacts with several pro-CO factors, may act as a master regulator by coordinating all the CO-promoting steps. Furthermore, our study suggests that CO distribution is regulated at all the different prophase I sub-stages through HEIP1 activity, thereby providing new insights into the orchestration of CO implementation and interference.

## Materials and Methods

*Heip1^-/-^* mutant were generated at the mice Transgenic Facility in Curie Institute, Paris (*SI Appendix* for details). All animal experiments were carried out according to the CNRS guidelines. Y2H, protein extracts and IP, RT-PCR assays, Histology and Immunocytology were performed using standard protocols. Details of antibody production, image acquisition, processing, and analysis are described in *SI Appendix*. Statistics were performed using Graphpad Prism v.10.4.2. Primers and antibodies are presented in Table S1 and S2. Counting of each experiment and replicate are presented in SI Dataset S01.

## Acknowledgments

We thank our lab members for helpful discussion and reading. We thank Fatima El Marjou and the Mice Transgenic Facility in Curie Institute for the generation of the *Heip1^-/-^*mouse model, Bernard de Massy and Christine Brun for chromosomes spreads from *Rec114^-/-^* mice, Attila Tóth for anti CNTD1 antibody and Bolcun-Filas lab for anti H1t antibody. We thank the Biocampus facilities from Montpellier for their service: the Réseau des Animaleries de Montpellier (RAM) for animal care, the Réseau d’Histologie Expérimentale de Montpellier (RHEM) for histology and Montpellier Resources Imagerie (MRI) for microscopy.

The Centre for Structural Biology (CBS) is a member of France-BioImaging (FBI) and the French Infrastructure for Integrated Structural Biology, two national infrastructures supported by the French National Research Agency (grant n. ANR-10-INBS-04-01 and ANR-10-INBS-05, respectively). TR’s group is supported by the CNRS INSERM ATIP-Avenir 2017 program, ANR CONDENSin3R (ANR-20-CE12-0016-02), ANR COMORE (ANR-24-CE12-2240-01) and La Ligue Contre le Cancer grants. LDT is funded by a PhD studentship from Université Montpellier, UM, ED CBS2. Work in V.B.’s group was supported by Fondation ARC and La Ligue Contre le Cancer grants. RM is supported by core funding from the Max Planck Society. This work was supported in part by NIH NICHD grant to Neil Hunter: R01HD109322. Neil Hunter is an Investigator of the Howard Hughes Medical Institute.

## References

1. N. Hunter, Meiotic Recombination: The Essence of Heredity. Cold Spring Harb Perspect Biol 7 (2015).

2. D. Zickler, N. Kleckner, Recombination, Pairing, and Synapsis of Homologs during Meiosis. Cold Spring Harb Perspect Biol 7 (2015).

3. B. de Massy, Initation of Meiotic Recombination: How and Where? Conversation and Specificities Among Eukaryotes. Annu Rev Genet (2013). 10.1146/annurev-genet-110711-155423.

4. T. Robert, N. Vrielynck, C. Mezard, B. de Massy, M. Grelon, A new light on the meiotic DSB catalytic complex. Semin Cell Dev Biol (2016). 10.1016/j.semcdb.2016.02.025.

5. T. Robert, et al., The TopoVIB-Like protein family is required for meiotic DNA double-strand break formation. Science 351, 943–9 (2016).

6. N. Vrielynck, et al., A DNA topoisomerase VI-like complex initiates meiotic recombination. Science 351, 939–43 (2016).

7. A. Pyatnitskaya, V. Borde, A. De Muyt, Crossing and zipping: molecular duties of the ZMM proteins in meiosis. Chromosoma 128, 181–198 (2019).

8. E. Cannavo, et al., Regulation of the MLH1-MLH3 endonuclease in meiosis. Nature 586, 618– 622 (2020).

9. H. Guillon, F. Baudat, C. Grey, R. M. Liskay, B. de Massy, Crossover and noncrossover pathways in mouse meiosis. Mol Cell 20, 563–73 (2005).

10. D. S. Kulkarni, et al., PCNA activates the MutLγ endonuclease to promote meiotic crossing over. Nature 586, 623–627 (2020).

11. K. Zakharyevich, S. Tang, Y. Ma, N. Hunter, Delineation of joint molecule resolution pathways in meiosis identifies a crossover-specific resolvase. Cell 149, 334–47 (2012).

12. S. M. Baker, et al., Involvement of mouse Mlh1 in DNA mismatch repair and meiotic crossing over. Nat Genet 13, 336–342 (1996).

13. N. K. Kolas, et al., Mutant meiotic chromosome core components in mice can cause apparent sexual dimorphic endpoints at prophase or X-Y defective male-specific sterility. Chromosoma 114, 92–102 (2005).

14. S. M. Lipkin, et al., Meiotic arrest and aneuploidy in MLH3-deficient mice. Nat Genet 31, 385– 390 (2002).

15. E. Marcon, P. Moens, MLH1p and MLH3p localize to precociously induced chiasmata of okadaic-acid-treated mouse spermatocytes. Genetics 165, 2283–2287 (2003).

16. P. B. Moens, et al., The time course and chromosomal localization of recombination-related proteins at meiosis in the mouse are compatible with models that can resolve the early DNA-DNA interactions without reciprocal recombination. J. Cell. Sci. 115, 1611–1622 (2002).

17. W. Edelmann, et al., Meiotic pachytene arrest in MLH1-deficient mice. Cell 85, 1125–1134 (1996).

18. A. Svetlanov, F. Baudat, P. E. Cohen, B. de Massy, Distinct functions of MLH3 at recombination hot spots in the mouse. Genetics 178, 1937–45 (2008).

19. A. De Muyt, et al., BLM helicase ortholog Sgs1 is a central regulator of meiotic recombination intermediate metabolism. Mol Cell 46, 43–53 (2012).

20. S. Tang, M. K. Wu, R. Zhang, N. Hunter, Pervasive and essential roles of the top3-rmi1 decatenase orchestrate recombination and facilitate chromosome segregation in meiosis. Molecular cell 57, 607–21 (2015).

21. D. Zickler, N. Kleckner, A Few of Our Favorite Things: Pairing, the Bouquet, Crossover Interference and Evolution of Meiosis. Semin Cell Dev Biol (2016). 10.1016/j.semcdb.2016.02.024.

22. D. Zickler, N. Kleckner, Meiosis: Dances Between Homologs. Annu Rev Genet 57, 1–63 (2023).

23. L. Chelysheva, et al., The Arabidopsis HEI10 is a new ZMM protein related to Zip3. PLoS Genet 8, e1002799 (2012).

24. K. Wang, et al., The role of rice HEI10 in the formation of meiotic crossovers. PLoS Genet 8, e1002809 (2012).

25. A. De Muyt, et al., E3 ligase Hei10: a multifaceted structure-based signaling molecule with roles within and beyond meiosis. Genes & development 28, 1111–23 (2014).

26. H. Qiao, et al., Antagonistic roles of ubiquitin ligase HEI10 and SUMO ligase RNF212 regulate meiotic recombination. Nat Genet 46, 194–199 (2014).

27. R. Yokoo, et al., COSA-1 reveals robust homeostasis and separable licensing and reinforcement steps governing meiotic crossovers. Cell 149, 75–87 (2012).

28. S. Durand, et al., Joint control of meiotic crossover patterning by the synaptonemal complex and HEI10 dosage. Nat Commun 13, 5999 (2022).

29. C. Girard, D. Zwicker, R. Mercier, The regulation of meiotic crossover distribution: a coarse solution to a century-old mystery? Biochem Soc Trans 51, 1179–1190 (2023).

30. C. Morgan, et al., Diffusion-mediated HEI10 coarsening can explain meiotic crossover positioning in Arabidopsis. Nat Commun 12, 4674 (2021).

31. Y. Li, et al., HEIP1 regulates crossover formation during meiosis in rice. Proc Natl Acad Sci U S A 115, 10810–10815 (2018).

32. D. K. Singh, et al., HEIP1 is required for efficient meiotic crossover implementation and is conserved from plants to humans. Proc Natl Acad Sci U S A 120, e2221746120 (2023).

33. Y. B. Condezo, et al., RNF212B E3 ligase is essential for crossover designation and maturation during male and female meiosis in the mouse. Proc Natl Acad Sci U S A 121, e2320995121 (2024).

34. M. Ito, et al., Distinct and interdependent functions of three RING proteins regulate recombination during mammalian meiosis. Proc Natl Acad Sci U S A 122, e2412961121 (2025).

35. J. O. Ward, et al., Mutation in mouse hei10, an e3 ubiquitin ligase, disrupts meiotic crossing over. PLoS Genet 3, e139 (2007).

36. L. Zhang, Z. Liang, J. Hutchinson, N. Kleckner, Crossover patterning by the beam-film model: analysis and implications. PLoS genetics 10, e1004042 (2014).

37. S. Fan, et al., A novel recombination protein C12ORF40/REDIC1 is required for meiotic crossover formation. Cell Discov 9, 1–20 (2023).

38. C. Tu, et al., Loss-of-function variants in human C12orf40 cause male infertility by blocking meiotic progression. Cell Discov 9, 87 (2023).

39. A. Reynolds, et al., RNF212 is a dosage-sensitive regulator of crossing-over during mammalian meiosis. Nat Genet 45, 269–78 (2013).

40. M. Barchi, et al., Surveillance of different recombination defects in mouse spermatocytes yields distinct responses despite elimination at an identical developmental stage. Mol Cell Biol 25, 7203–15 (2005).

41. F. A. T. de Vries, et al., Mouse Sycp1 functions in synaptonemal complex assembly, meiotic recombination, and XY body formation. Genes Dev. 19, 1376–1389 (2005).

42. A. Bondarieva, et al., Proline-rich protein PRR19 functions with cyclin-like CNTD1 to promote meiotic crossing over in mouse. Nat Commun 11, 3101 (2020).

43. R. Kumar, H. M. Bourbon, B. de Massy, Functional conservation of Mei4 for meiotic DNA double-strand break formation from yeasts to mice. Genes Dev 24, 1266–80 (2010).

44. J. Ribeiro, E. Abby, G. Livera, E. Martini, RPA homologs and ssDNA processing during meiotic recombination. Chromosoma 125, 265–276 (2016).

45. A. De Muyt, et al., A meiotic XPF-ERCC1-like complex recognizes joint molecule recombination intermediates to promote crossover formation. Genes Dev. 32, 283–296 (2018).

46. Y. Li, et al., M1AP interacts with the mammalian ZZS complex and promotes male meiotic recombination. EMBO Rep 24, e55778 (2023).

47. L. Jessop, B. Rockmill, G. S. Roeder, M. Lichten, Meiotic chromosome synapsis-promoting proteins antagonize the anti-crossover activity of sgs1. PLoS Genet 2, e155 (2006).

48. S. D. Oh, et al., BLM ortholog, Sgs1, prevents aberrant crossing-over by suppressing formation of multichromatid joint molecules. Cell 130, 259–72 (2007).

49. F. Baudat, K. Manova, J. P. Yuen, M. Jasin, S. Keeney, Chromosome synapsis defects and sexually dimorphic meiotic progression in mice lacking Spo11. Mol Cell 6, 989–98 (2000).

50. R. Kumar, et al., Mouse REC114 is essential for meiotic DNA double-strand break formation and forms a complex with MEI4. Life Sci Alliance 1, e201800259 (2018).

51. M. Stanzione, et al., Meiotic DNA break formation requires the unsynapsed chromosome axis-binding protein IHO1 (CCDC36) in mice. Nat Cell Biol (2016). 10.1038/ncb3417.

52. J. K. Holloway, X. Sun, R. Yokoo, A. M. Villeneuve, P. E. Cohen, Mammalian CNTD1 is critical for meiotic crossover maturation and deselection of excess precrossover sites. J Cell Biol 205, 633–641 (2014).

53. B. Kneitz, et al., MutS homolog 4 localization to meiotic chromosomes is required for chromosome pairing during meiosis in male and female mice. Genes Dev 14, 1085–1097 (2000).

54. Q. Zhang, J. Shao, H.-Y. Fan, C. Yu, Evolutionarily-conserved MZIP2 is essential for crossover formation in mammalian meiosis. Commun Biol 1, 147 (2018).

55. Q. Zhang, S.-Y. Ji, K. Busayavalasa, C. Yu, SPO16 binds SHOC1 to promote homologous recombination and crossing-over in meiotic prophase I. Sci Adv 5, eaau9780 (2019).

56. M. F. Guiraldelli, C. Eyster, J. L. Wilkerson, M. E. Dresser, R. J. Pezza, Mouse HFM1/Mer3 is required for crossover formation and complete synapsis of homologous chromosomes during meiosis. PLoS Genet 9, e1003383 (2013).

57. D. C. Nageswaran, et al., HIGH CROSSOVER RATE1 encodes PROTEIN PHOSPHATASE X1 and restricts meiotic crossovers in Arabidopsis. Nat Plants 7, 452–467 (2021).

58. Z. Chang, et al., The plant-specific ABERRANT GAMETOGENESIS 1 gene is essential for meiosis in rice. J Exp Bot 71, 204–218 (2020).

59. M. Biot, et al., Principles of chromosome organization for meiotic recombination. Mol Cell 84, 1826–1841.e5 (2024).

60. N. R. Bhagwat, et al., SUMO is a pervasive regulator of meiosis. Elife 10, e57720 (2021).

61. H. B. Rao, et al., A SUMO-ubiquitin relay recruits proteasomes to chromosome axes to regulate meiotic recombination. Science (2017). 10.1126/science.aaf6407.

62. C. S. Eichinger, S. Jentsch, Synaptonemal complex formation and meiotic checkpoint signaling are linked to the lateral element protein Red1. Proc Natl Acad Sci U S A 107, 11370– 11375 (2010).

63. J. H. Joo, et al., RPA interacts with Rad52 to promote meiotic crossover and noncrossover recombination. Nucleic Acids Research 52, 3794–3809 (2024).

64. S. Lahiri, L. Y, H. Mm, M. I, MutSγ-Induced DNA Conformational Changes Provide Insights into Its Role in Meiotic Recombination. Biophysical journal 115 (2018).

65. T. Snowden, S. Acharya, C. Butz, M. Berardini, R. Fishel, hMSH4-hMSH5 recognizes Holliday Junctions and forms a meiosis-specific sliding clamp that embraces homologous chromosomes. Mol Cell 15, 437–451 (2004).

66. M. Arai, S. Suetaka, K. Ooka, Dynamics and interactions of intrinsically disordered proteins. Curr Opin Struct Biol 84, 102734 (2024).

67. E. L. Sipko, G. F. Chappell, R. B. Berlow, Multivalency emerges as a common feature of intrinsically disordered protein interactions. Curr Opin Struct Biol 84, 102742 (2024).

68. M. F. Guiraldelli, et al., SHOC1 is a ERCC4-(HhH)2-like protein, integral to the formation of crossover recombination intermediates during mammalian meiosis. PLoS Genet 14, e1007381 (2018).

69. H. Kaur, A. De Muyt, M. Lichten, Top3-rmi1 DNA single-strand decatenase is integral to the formation and resolution of meiotic recombination intermediates. Molecular cell 57, 583–94 (2015).

70. A. H. Peters, A. W. Plug, M. J. van Vugt, P. de Boer, A drying-down technique for the spreading of mammalian meiocytes from the male and female germline. Chromosome Res 5, 66–8 (1997).

71. C. Grey, F. Baudat, B. de Massy, Genome-wide control of the distribution of meiotic recombination. PLoS Biol 7, e35 (2009).

72. J. Cau, L. D. Toe, A. Zainu, F. Baudat, T. Robert, “MeiQuant”: An Integrated Tool for Analyzing Meiotic Prophase I Spread Images. Methods Mol Biol 2770, 263–285 (2024).

