## Appendix for "HEIP1 orchestrates pro-crossover protein activity during mammalian meiosis"

### Supplementary Figures

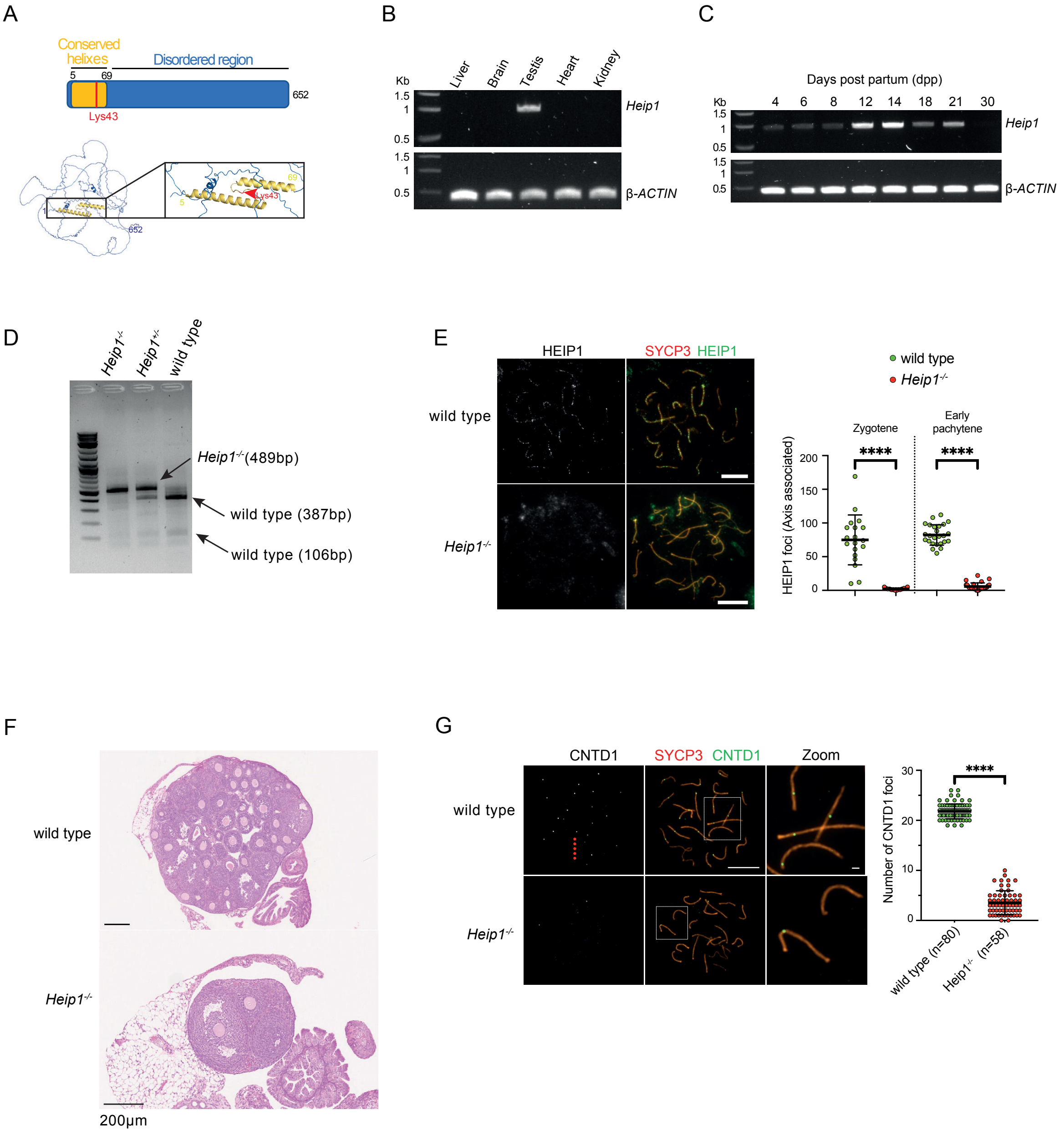


### Figure S1

**Figure S1:**
(A) (top) Schematic and aphafold2 (bottom) representation of HEIP1 protein. Alpha helixes are shown in yellow while the disorder region is shown in blue. The mutation created in the Lys 43 is represented in red. (B and C) Levels of *Heip1* mRNA in multiple mouse tissues. β-Actin served as the loading control. (D) *Heip1* genomic fragments were amplified with Heip1-P5 and Heip1-P6 primers followed by a Hpy188III digestion. *Heip1*$^{-/-}$ has lost a Hpy188III restriction site leading to a PCR fragment of 489 bp instead of two wild type bands of 387 bp and 106 bp. (E) co-immunolocalization of SYCP3 and HEIP1 in wild type and *Heip1*$^{-/-}$ spermatocytes during late zygotene and early pachytene stage. Plot at the right of the panel show the quantitation of HEIP1 foci; ****P < 0.0001 (two-tailed unpaired Mann-Whitney test). Scale bar, 10 µm. (F) Hematoxylin and eosin staining of ovary sections from wild type and *Heip1*$^{-/-}$ at 21 dpp. Scale bar, 200 µm. (G) Immunostaining of CNTD1 (green) and SYCP3 (Orange) on chromosome spreads from wild type and *Heip1*$^{-/-}$ spermatocytes at 18 dpp. White squares correspond to the magnified images on the right. The number of MLH1 foci is presented on the right. Scale bar, 10 µm (1um for the magnified image). For all graphs, ****P < 0.0001 (two-tailed unpaired Mann-Whitney test).

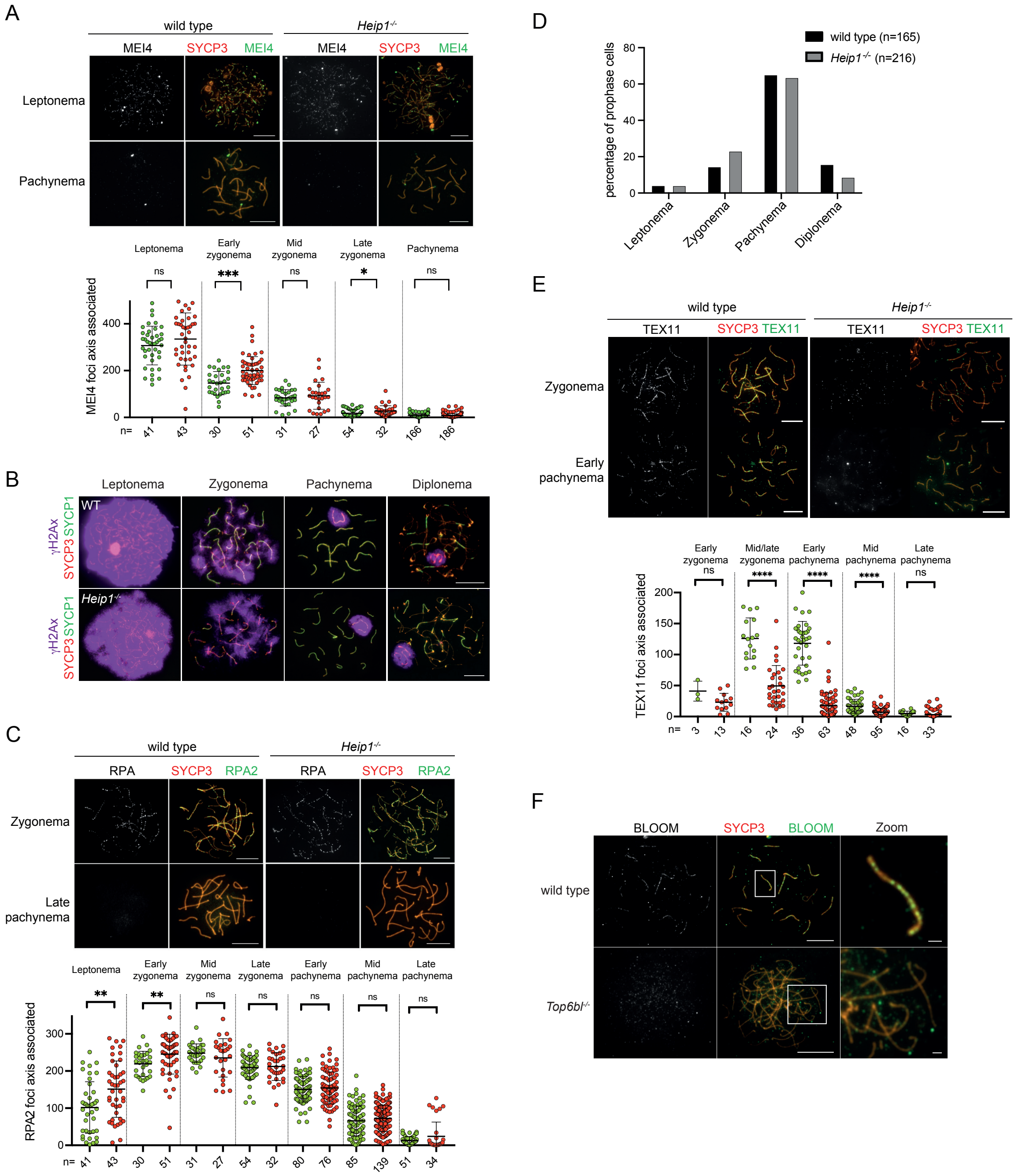
A
wild type
Heip1-/-
MEI4
SYCP3 MEI4
Leptonema
Pachynema
Leptonema
Early zygonema
Mid zygonema
Late zygonema
Pachynema
ns
***
ns
*
ns
MEI4 foci axis associated
n=
41
43
30
51
31
27
54
32
166
186
B
Leptonema
Zygonema
Pachynema
Diplonema
γH2Ax SYCP3 SYCP1
WT
Heip1-/-
C
wild type
Heip1-/-
RPA
SYCP3 RPA2
Zygonema
Late pachynema
Leptonema
Early zygonema
Mid zygonema
Late zygonema
Early pachynema
Mid pachynema
Late pachynema
**
**
ns
RPA2 foci axis associated
n=
41
43
30
51
31
27
54
32
80
76
85
139
51
34
D
wild type (n=165)
Heip1-/- (n=216)
percentage of prophase cells
Leptonema
Zygonema
Pachynema
Diplonema
E
wild type
Heip1-/-
TEX11
SYCP3 TEX11
Zygonema
Early pachynema
Early zygonema
Mid/late zygonema
Early pachynema
Mid pachynema
Late pachynema
ns
****
****
****
ns
TEX11 foci axis associated
n=
3
13
16
24
36
63
48
95
16
33
F
BLOOM
SYCP3 BLOOM
Zoom
wild type
Top6bl-/-

**Figure S2**

**Figure S2:**
(A) Top: Immunostaining of MEI4 (green) and SYCP3 (orange) on chromosome spreads of 18 dpp wild type and *Heip1*$^{-/-}$ spermatocytes at different prophase I substages. Scale bar, 10 µm Bottom: quantitation of MEI4 foci; ns, not significant, *P < 0.1, ***P < 0.001 (two-tailed unpaired Mann-Whitney test). The number of nuclei used for the quantification is shown. (B) Immunostaining of $\gamma$H2AX (purple), SYCP3 (orange) and SYCP1 (green) on chromosome spreads of 18 dpp wild type and *Heip1*$^{-/-}$ spermatocytes at different prophase I substages. Scale bar, 10 µm. (C) Top: Immunostaining of RPA2 (green) and SYCP3 (orange) on chromosome spreads of 18 dpp wild type and *Heip1*$^{-/-}$ spermatocytes at different prophase I substages. Scale bar, 10 µm Bottom: quantitation of MEI4 foci; ns, not significant, **P < 0.01 (two-tailed unpaired Mann-Whitney test). The number of nuclei used for the quantification is shown. (D) Prophase I staging of wild type and *Heip1*$^{-/-}$ of 18 dpp spermatocytes. The number of nuclei used for the quantification is shown. (E) Top: Immunostaining of TEX11 (green) and SYCP3 (orange) on chromosome spreads of 60 dpp wild type and *Heip1*$^{-/-}$ spermatocytes at different prophase I substages. Scale bar, 10 µm Bottom: quantitation of MEI4 foci; ns, not significant, ****P < 0.0001 (two-tailed unpaired Mann-Whitney test). The number of nuclei used for the quantification is shown. (F) Immunostaining of BLOOM (green) and SYCP3 (orange) on chromosome spreads of 18 dpp wild type and *Top6bl*$^{-/-}$ spermatocytes at different prophase I substages. In the *Top6bl*$^{-/-}$ mutant, in which DSBs do not form, Bloom foci were almost lost showing that they strictly depend on recombination. Scale bar, 10 µm

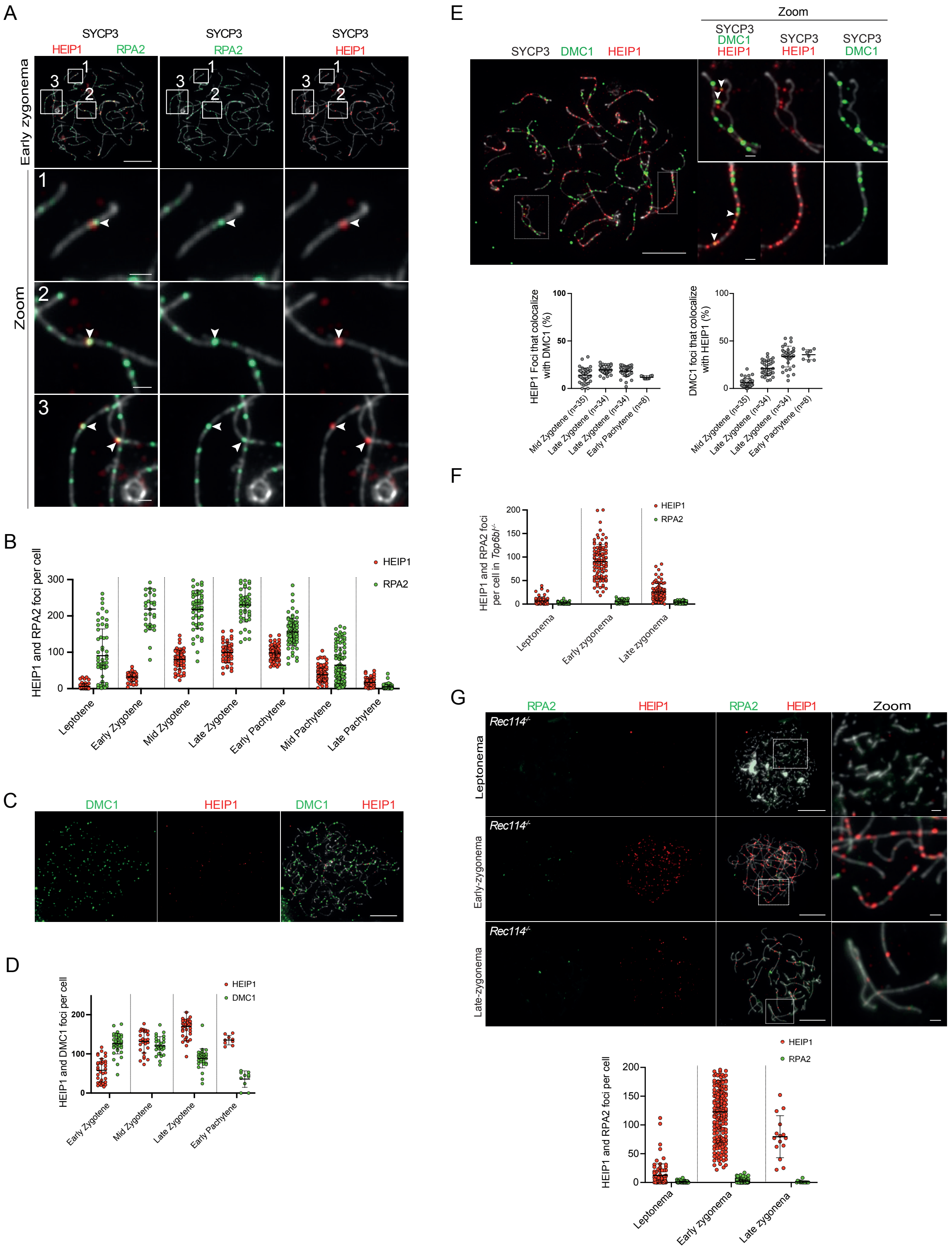
A
SYCP3
HEIP1
RPA2
Early zygonema
Zoom
B
HEIP1 and RPA2 foci per cell
HEIP1
RPA2
Leptotene
Early Zygotene
Mid Zygotene
Late Zygotene
Early Pachytene
Mid Pachytene
Late Pachytene
C
DMC1
HEIP1
D
HEIP1 and DMC1 foci per cell
Early Zygotene
Mid Zygotene
Late Zygotene
Early Pachytene
E
Zoom
SYCP3
DMC1
HEIP1
HEIP1 Foci that colocalize with DMC1 (%)
DMC1 foci that colocalize with HEIP1 (%)
Mid Zygotene (n=35)
Late Zygotene (n=34)
Late Zygotene (n=34)
Early Pachytene (n=8)
F
HEIP1 and RPA2 foci per cell in Top6bl-/-
Leptonema
Early zygonema
Late zygonema
G
Rec114-/-
Leptonema
Early-zygonema
Late-zygonema
Zoom
HEIP1 and RPA2 foci per cell

**Figure S3**

**Figure S3:**

(A) Co-immunolocalization of HEIP1 (red) and RPA2 (green) during early zygotene stage of wild type spermatocytes. Three regions of the nucleus are shown in the enlarged image 1 to 3 from the white square. Arrowheads indicate the co-localization between HEIP1 and RPA2 at the chromosome axes stained by SYCP3 (grey). Scale bar, 10 µm (1um for the magnified image). (B) Quantification of HEIP1 (red) and RPA2 (green) foci during prophase I of wild type spermatocytes. (C) Co-immunolocalization of HEIP1 (red) and DMC1 (green) at leptotene stage of wild type spermatocytes. Scale bar, 10 µm (D) Quantification of HEIP1 (red) and DMC1 (green) foci during early stages of prophase I of wild type spermatocytes. (E) Chromosome spreads from 18 dpp wild type prophase I spermatocyte nuclei immunostained for SYCP3 (grey), HEIP1 (red) and DMC1 (green). Magnified images show representative chromosomes. Scale bars, 10 µm for full nuclei and 1 µm for magnified images. Arrowheads indicate co-localized HEIP1 and DMC1 foci. The plots at the bottom show the percentage of co-localization between HEIP1 and DMC1 at early stage of prophase I. The number of nuclei used for the quantification is shown. (F) Quantitation of RPA2 and HEIP1 foci at different stages of *Top6bl*$^{-/-}$ prophase I. Bars indicate the mean ± SD. Quantification is related to the images presented in Figure 3E. (G) Chromosome spreads from 14 dpp *Rec114*$^{-/-}$ prophase I spermatocyte nuclei immunostained for SYCP3 (grey), HEIP1 (red) and RPA2 (green). Magnified images show representative chromosomes. Scale bars, 10 µm for full nuclei and 1 µm for magnified images. Plot at the bottom of the panel show the quantitation of RPA2 and HEIP1 foci at different stages of *Rec114*$^{-/-}$ prophase I. Bars indicate the mean ± SD.

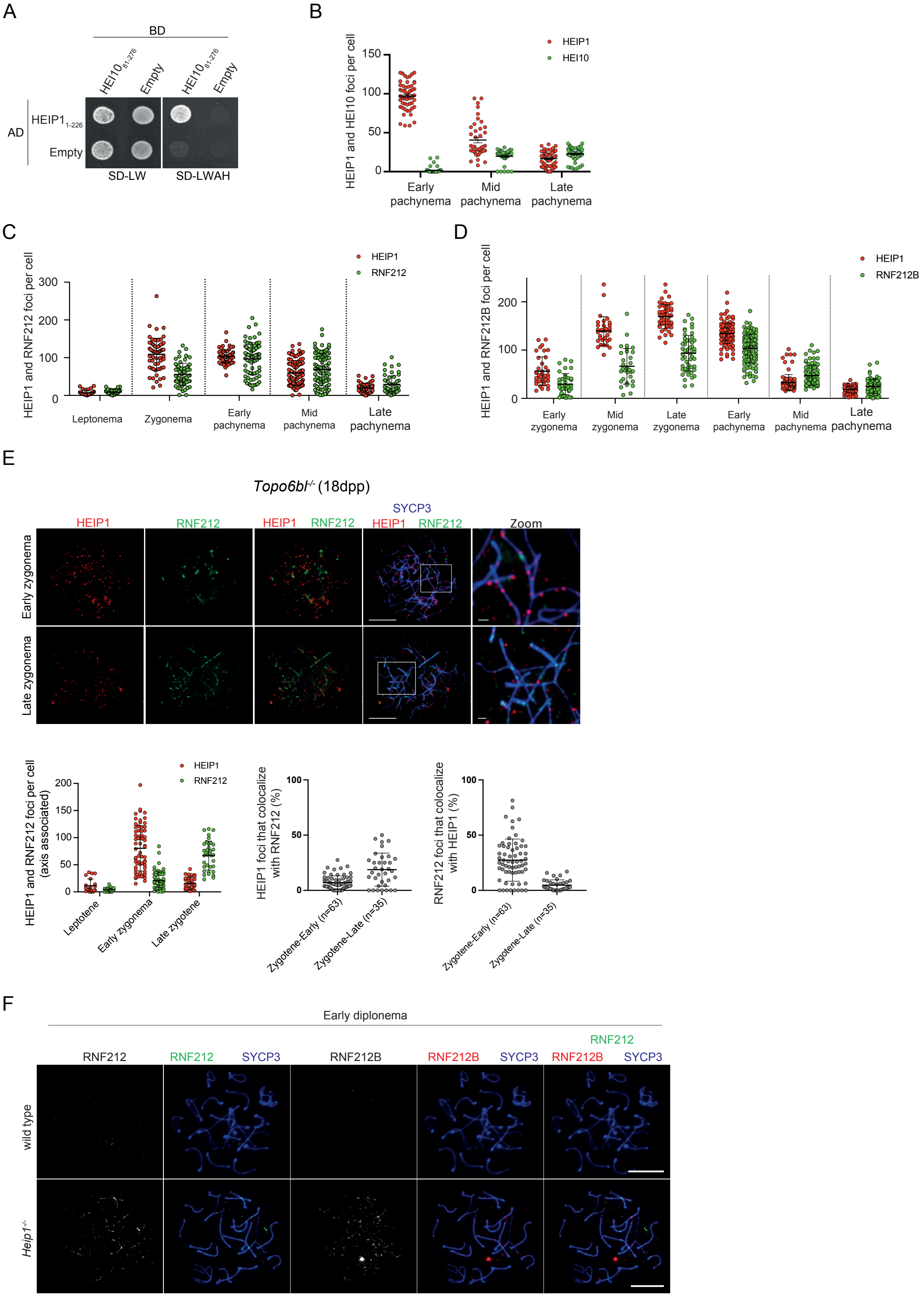

A
BD
HEI10 81-276
Empty
HEI10 81-276
Empty
AD
HEIP1 1-226
Empty
SD-LW
SD-LWAH
B
HEIP1 and HEI10 foci per cell
HEIP1
HEI10
Early pachynema
Mid pachynema
Late pachynema
C
HEIP1 and RNF212 foci per cell
HEIP1
RNF212
Leptonema
Zygonema
Early pachynema
Mid pachynema
Late pachynema
D
HEIP1 and RNF212B foci per cell
HEIP1
RNF212B
Early zygonema
Mid zygonema
Late zygonema
Early pachynema
Mid pachynema
Late pachynema
E
Topo6bl-/- (18dpp)
HEIP1
RNF212
SYCP3
Zoom
Early zygonema
Late zygonema
HEIP1 and RNF212 foci per cell (axis associated)
Leptotene
Early zygonema
Late zygotene
HEIP1 foci that colocalize with RNF212 (%)
Zygotene-Early (n=63)
Zygotene-Late (n=35)
RNF212 foci that colocalize with HEIP1 (%)
F
Early diplonema
RNF212
RNF212B
SYCP3
wild type
Heip1-/-


**Figure S4**

**Figure S4:**
(A) Yeast two-hybrid assay showing interaction between, the N-terminal part of HEIP1 (HEIP11-226) and the C-terminal part of HEI10 (HEI1081-276). Yeast cells expressing Gal4 activation (AD) and binding (BD) domain fused to the indicated proteins, or empty vectors as controls, were spotted. (B) Quantification of the HEIP1 (red) and HEI10 (green) foci numbers during pachytene substages. (C) Quantification of the HEIP1 (red) and RNF212 (green) foci numbers at different prophase I substages. (D) Quantification of HEIP1 and RNF212B at different prophase I substages. (E) Immunostaining of SYCP3 (blue), HEIP1 (Red) and RNF212 (green) from 18 dpp spermatocytes at different prophase I substages of *Topo6bl*$^{-/-}$. (F) Immunostaining of SYCP3 (blue), RNF212 (green) and RNF212B (red) from 60 dpp wild type spermatocytes at early diplonema.

**Supplementary Material and Methods**

**Mouse strains.** Mouse strains were in the C57BL/6 J background. Housing conditions were: 22 °C, 55% humidity, 12h/12h dark/light cycle (8 am/8 pm in summer and 7 am/7 pm in winter). All animal experiments were carried out according to the CNRS guidelines.

**Generation of mutant mice by CRISPR/Cas9.** Mutant mice were created at the Jackson Laboratory using the CRISPRCas9 technology with two different guides and one donor oligonucleotide (Table S1). Guides were selected to minimize off target effects. The *Heip1*$^{-/-}$ allele is the result of non-homologous repair and has a 4 bp deletion (Figure 1). Founders were backcrossed with C57BL/6J mice to obtain heterozygous animals. The predicted HEIP1 protein expressed from the mutant allele and the genotyping strategies are shown in Figure S1A and D.

**Yeast two-hybrid assay.** Full-length mouse *Heip1* and *Hei10* and their truncated versions were PCR-amplified from mouse testis cDNA (see Table S1 for primers). PCR products were cloned in plasmids derived from the two hybrid vectors pGADT7 (GAL4-activating domain) and pGBKT7 (GAL4-binding domain), creating N-terminal fusions and transformed in the yeast haploid strains Y187 and AH109 (Clontech). Yeast two-hybrid assays were performed and interactions were scored on selective media exactly as described in (45). An interaction was defined compared to the growth seen in the negative control (*i.e.* the combination between the GAL4BD-bait protein in the presence of GAL4AD-only; 'empty' on the Figures). Any combination that grows better than this control on the selective media was considered as an interaction.

**Preparation of mouse protein extracts, co-immunoprecipitation and western blotting.** Whole cell protein extracts were prepared from four frozen testes/genotype collected at 18 dpp. After protein extraction by homogenizing cells with a Dounce homogenizer in HNTG buffer [150 mM NaCl, 20 mM HEPES pH7.5, 1% Triton X-100, 10% glycerol, 1 mM MgCl, Complete protease Inhibitor (Roche 11873580001)], followed by sonication, benzonase (250 U) was added at 4 °C for 1 h. After centrifugation (16000 g, 4 °C, 10 min) to remove debris, immunoprecipitation was performed with 5 µg of homemade anti-HEIP1 antibody. For each immunoprecipitation, 3.5 mg of whole cell protein extract and 50 µl of Protein A Dynabeads (Invitrogen 10001D) were used. Then, immunoprecipitates were resuspended in 40 µl of Laemmli buffer and HEIP1 immunoprecipitation was assessed by western blotting using antibodies presented in Table S2.

**Antibody production.** <u>HEI10 antibody:</u> Polyclonal antibodies against HEI10 were raised in guinea pigs. Full length *Hei10* was PCR-amplified from mouse testis cDNA using the primer presented in Table S1. The *Hei10* PCR fragment was cloned using the In-Fusion cloning method into a modified pDB vector carrying a 3C protease cleavable *6*His + thioredoxin (Trx) tag. *E. coli* (DE3) cells (Agilent Technologies Inc.) transformed with the 6His-Trx-3C-Hei10 expression vector were grown in 2L of LB at 37°C to OD600 = 0.8, and protein expression was induced by adding 0.5 mM IPTG at 37°C for 3h.
<u>HEIP1 antibody:</u> Polyclonal antibodies against mouse HEIP1 were raised in rabbits. Truncated (484-652) and full length *Heip1* were PCR-amplified from mouse testis cDNA using the primers presented in Table S1. The Heip1 (484-652) PCR fragment was subcloned using the Gateway cloning system into the pDONR207 vector (Invitrogen), leading to pENTR207-Heip1 (484-652). From pENTR207-Heip1 (484-652), the Heip1 (484-652) PCR fragment was transferred by LR recombination (Invitrogen) into the pDEST566 vector carrying a TEV protease cleavable 6His + Maltose Binding Protein (MBP) tag. Transformed *E. coli* (DE3) cells were grown in 2L of LB at 37°C to OD600 = 0.8, and protein expression was induced with 0.5 mM IPTG followed by incubation at 37°C for 3h.

**RT-PCR assays.** Total RNA was extracted from 18dpp testes (unless indicated) with the miRNeasy Mini Kit (Qiagen) according to the manufacturer's instructions. For RT-PCR, first-strand DNA was synthesized using oligo d(T)18 (Ambion), SuperScriptIII (Invitrogen) and total RNA (1-2 µg). The open reading frames of *Heip1* and *Hei10* were amplified using standard PCR conditions and the primer pairs in Table S1. PCR cycling conditions were: 3 min at 94 °C, 35 cycles of 30 sec at 94 °C, 30 sec at 60 °C, and 2 min or 1 min at 72 °C, followed by 5 min at 72 °C.

**Histological analysis of paraffin sections.** Mouse testes and ovaries were fixed respectively in PFA and Bouin's solution. Testes and ovaries were embedded in paraffin and cut in 3µm-thick sections followed by hematoxylin-eosin (HE) staining. Sections were scanned using the automated tissue slide-scanning tool of a Hamamatsu NanoZoomer Digital Pathology system.

**Immunocytology.** Spermatocyte spreads were prepared using the dry down technique, as described in (70). Briefly, a suspension of testis cells was prepared in PBS, and then incubated in hypotonic solution at room temperature for 8 min. Cells were centrifuged, resuspended in 66 mM sucrose solution and spread on slides with 1% paraformaldehyde/0.05% Triton X-100. Slides were dried in a humid chamber for 1 to 2h. Immunostaining was performed using a milk-based blocking buffer (5% milk, 5% donkey serum in PBS) (71). Slides were incubated with primary antibodies at room temperature overnight and with secondary antibodies at 37 °C for 1h. Nuclei were stained with 2 µg/ml 4'−6-diamidino-2-phenylindole (DAPI) during the final washing step.
All image analyses (foci count, intensity measurement) were performed using Fiji/ImageJ 1.53t, with the "MeiQuant" set of tools(72) available on github (https://github.com/MontpellierRessourcesImagerie/meiosis_bar), to the exclusion of MLH1 and CNTD1 foci number count performed manually.
